# High complexity cellular barcoding and clonal tracing reveals stochastic and deterministic parameters of radiation resistance

**DOI:** 10.1101/2020.10.13.337519

**Authors:** Anne Wursthorn, Christian Schwager, Ina Kurth, Claudia Peitzsch, Christel Herold-Mende, Jürgen Debus, Amir Abdollahi, Ali Nowrouzi

## Abstract

The impact of functional heterogeneity in response to radiation therapy is poorly understood to the present. It remains elusive whether clonal selection of tumor cells in response to ionizing radiation (IR) is a deterministic or stochastic process. We applied high-resolution lentiviral cellular barcoding for quantitative clonal tracking and deconvolution of clonal dynamics in response to IR. Clonal tracking of over 400.000 HNSCC patient-derived tumor cells and the analyses of over 1500 million sequencing reads in clonogenic survival assays reveals that fractionated IR induced a strong selective pressure for clonal reduction. This significantly exceeded uniform clonal survival probabilities indicative for a strong clone-to clone difference within tumor cell lines. IR induced clonal reduction affected the majority of tumor cells ranging between 96-75% and correlated to the degree of radiation sensitivity. Survival and clonogenicity is characterized by an intensive clonal distortion and dominance of individual tumor cells. Survival to IR is driven by a deterministic clonal selection of a smaller population which commonly survives radiation, while increased clonogenic capacity is a result of clonal competition of cells which have been selected stochastically. A 2-fold increase in radiation resistance results in a 4-fold (p<0.05) higher deterministic clonal selection showing that the ratio of these parameters is amenable to radiation sensitivity which correlates to prognostic biomarkers of HNSCC. Evidence for the existence of a rare subpopulation with an intrinsically radiation resistant phenotype commonly surviving IR was found at a frequency of 0.6-3.3% (p<0.001, FDR 3%). With cellular barcoding we introduce a novel functional heterogeneity associated qualitative readout for tracking dynamics of clonogenic survival in response to radiation. This enables the quantification of intrinsically radiation resistant tumor cells from patient samples and reveals the contribution of stochastic and deterministic clonal selection processes in response to IR.

## Introduction

Functional heterogeneity within tumor cell populations remains a major challenge for understanding the cellular response to ionizing radiation (IR) and the development of resistance mechanisms to radiotherapy^1^. The clonogenic survival assay is considered as the gold standard to determine radiosensitivity of tumor cells^2^. The radiation sensitivity of tumor cell lines and patient-derived tumor cells determined in clonogenic assays correlates to the tumor control rates and overall clinical outcome of radiation therapy^3–7^. In clonogenic survival assays the indefinite proliferative capability of single cell-derived clones is measured by the quantification of cell colonies consisting of at least 50 cells which formed within 10-14 days^2^. The quantity of single cell clones that have clonogenic survival is used to determine the survival fraction (SF) of the radiation dose. The implicit assumption these studies take is that every cell clone of a cell population reacts in a uniform manner. However, it is becoming increasingly apparent that considerable phenotypic and functional heterogeneity in tumor cell and patient-derived tumor lines exists^8^. The impact of functional heterogeneity on the radiosensitivity is to our knowledge not dissected at single clone resolution.

An important aspect of radiation resistance that has not been addressed comprehensively is the underlying clonal dynamics of cells that survive radiation and show lower sensitivity to radiation. Tumor cells that display an increased probability to survive radiation may originate as the clonal progeny of a pre-existing cell population with distinct molecular features that promote survival in response to IR. On the other hand, genomic heterogeneity can be induced by IR itself through *de novo* DNA mutations. Such mutations may have an impact on functional heterogeneity potentially increasing the ability to survive IR through acquired mechanisms^9^. We reasoned that radiation survival of tumor cells may be driven on a deterministic route through the existence of clones that have intrinsic molecular features that promote the survival in response to radiation. In contrast, clonogenic radiation survival and resistance to radiation may also occur in a stochastic manner by the selection of random cell clones, which might have acquired molecular capacities to survive radiation through IR-induced *de novo* mutations or have not received a dose sufficient for cell killing. Insights into clonal dynamics and clonal survival in response to radiation have only begun to be partially addressed in the molecular follow up of clinical trials^10,11^. Although clinically highly important, survival probabilities of individual cells have not been addressed in the context of functional heterogeneity due to the lack of molecular high-throughput methods.

Clonal marking by tracing lentiviral integration sites has been applied on a variety of cells such as tumor cells, blood cells and hepatocytes^12–14^. Using integration site analyses of lentiviral vectors for clonal tracing however is challenging since genomic access of integration sites usually requires the use of a combination of restriction enzymes^15,16^ which can be overcome by using lentiviral barcode libraries^17,18^. With lentiviral high-complexity cellular barcoding and longitudinal tracing of barcoded tumor cells we here introduce an experimental approach, which enables to investigate clonal origins and clonal dynamics of radiation resistance in clonogenic assays. To dissect and compare clonal dynamics and survival probabilities in clonogenic assays in response to IR the common head and neck squamous cancer cell line FaDu and two patient-derived cancer lines, which differ in their radiation sensitivity and molecular characteristics, were barcoded and analyzed. The quantitative assessment of clone numbers and clonal abundance with next generation sequencing (NGS) of irradiated cell population provides evidence that survival and clonogenicity can be seen as two different functional steps in clonogenic assays that are driven by two different tumor cell populations which survive and clonally expand deterministically or stochastically. Survival is dominantly driven by a smaller tumor cell population which commonly survives exposure to radiation in a deterministic manner whereas clonogenicity is a result of clonal competition between clones that have survived radiation in a deterministic and stochastic manner. The ratio between the stochastic and deterministic survival probability is amenable to radiation sensitivity and the frequency of prognostic biomarkers in HNSCC providing an index for the frequency of radiation resistant tumor cell clones in patient-derived tumor samples. Lentiviral cellular barcoding in combination with clonogenic survival assays enables the quantitative assessment of clonal dynamics in response to irradiation based on NGS rather than manual counting of cell colonies. This adds a biological and clinically important value to commonly used methods in radiation oncology, by linking clonal identity to clonogenicity in high resolution.

## Results

### A phenotypic characterization of an established HNSCC cell line and patient-derived HNSCC cultures reveals differences in the frequency in biomarkers of radiation resistance

For clonal marking and subsequent clonal tracing of individual tumor cells in response to ionizing radiation (IR) the HNSCC cell line FaDu and two previously described patient-derived HNSCC cell lines^19,20^ were used. One patient line was derived from a primary tumor (HNO206) and one from lymph node metastasis (HNO407). The HNSCC lines used were first molecularly characterized. RNA sequencing (RNA-seq) was performed on FaDu, HNO206 and HNO407 to compare gene expression patterns between the HNSCC cell lines. Spearman correlation analysis combined with hierarchical clustering (Manhattan distance) revealed that gene expression patterns for each HNSCC line is unique and that the HNO206 line clustered separately from the FaDu and HNO407 line (**Fig. 1A**). In total, the expression of 1587 genes were differentially regulated in HNSCC cell lines and affected cellular processes regulating cellcell communication, metabolism of lipids and RNA, senescence and extracellular matrix interactions (p<0.00005, ANOVA **Supplementary Fig. 1A, Supplementary Table 1A, Supplementary Data 1A**). We further analyzed the gene expression differences between the HNO407 compared to HNO206 line (**Fig. 1B**) resulting in a total of 464 genes (**Supplementary Data 1B**), which were significantly differentially regulated more than 4-fold (p<0.001, unpaired t-test). The differentially regulated genes were mainly involved in functional categories affecting extracellular matrix interaction, collagen synthesis, and integrin interactions comprising cellular processes of epithelial to mesenchymal transition (EMT) (**Supplementary Table 1B**). Gene Set Enrichment Analysis (GSEA) verified the differences in the upregulation of EMT regulation in the HNO407 (p<0.05, multiple hypothesis testing, **Fig. 1C, Supplementary Table 1C**). EMT is a driver of tumor progression and metastasis and has been proposed to play a role in radioresistance^21,22^. Therefore, we evaluated the expression of individual genes involved in the regulation of EMT^23^ and found the mesenchymal markers vimentin (VIM), integrin α-5 (ITGA5) and N-cadherin (CDH2) as well as the EMT regulator transforming growth factor (TGF)-b-induced (TGFBI) is upregulated in HNO407 (p<0.001, unpaired t-test, **Fig. 2A**). Furthermore, genes which are considered as prognostic factors to distinguish good versus bad clinical outcome in HNSCC patients (ABCG2, CD44, CDKN1A, EGFR, SOX9) are upregulated in HNO407, whereas other markers (AKT2, FOS, PTEN) including the radioresistance-associated biomarker ALDH (ALDH1A3) were downregulated in HNO407 compared to HNO206 (**Fig. 2A**).

**Figure 1 -.**
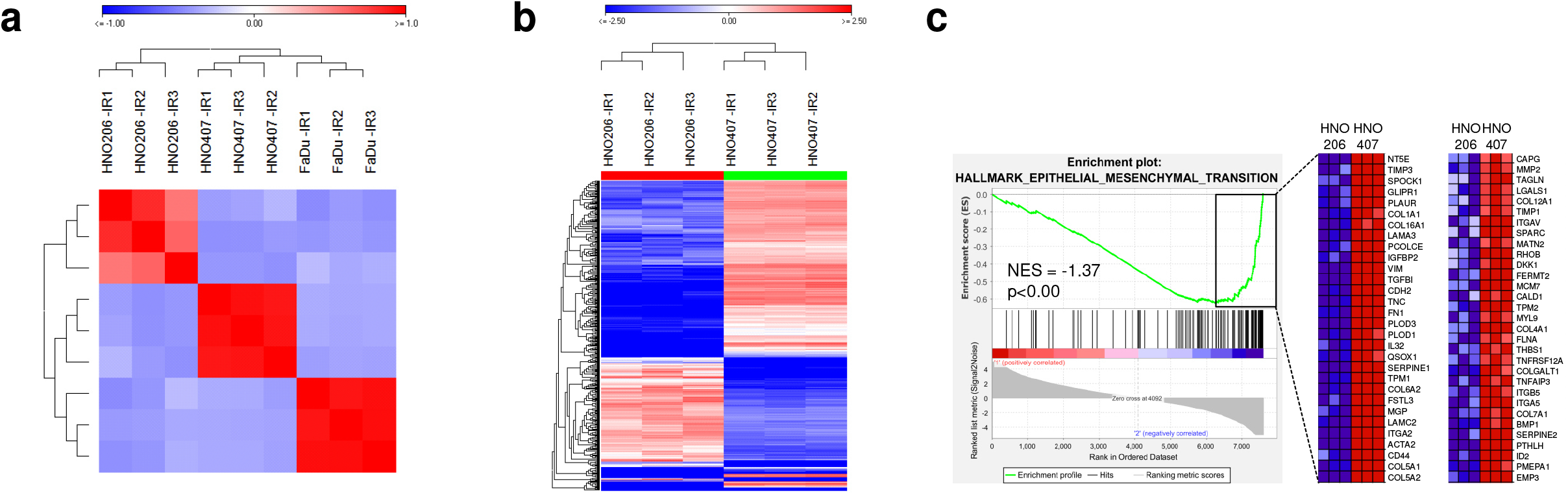
Transcriptional characterization of the HNSCC lines reveals gene expression signatures relevant for accessing the radiosensitivity. A Spearman correlation analysis of RNA-seq data demonstrated unique gene expression patterns for HNSCC cell lines and hierarchical clustering (Manhattan distance) revealed separated clustering of HNO206 from FaDu and HNO407. B Heatmap of >4-fold differentially expressed genes in the patient-derived cell lines HNO206 (derived from a primary tumor) compared to HNO407 (derived from a lymph node metastasis), (unpaired t-test, p<0.001, log_2_). C Regulation of Gene Set Enrichment Analysis (GSEA) of HNO206 versus HNO407 using the Hallmark gene set collection from the Molecular Signature Database (MSigDB) (multiple hypothesis testing, p<0.05, NES = normalized enrichment score). Heatmap shows the relative level of gene expression (red = high, blue = low) for the most significant genes that contributed to the hallmark.

**Figure 2 -.**
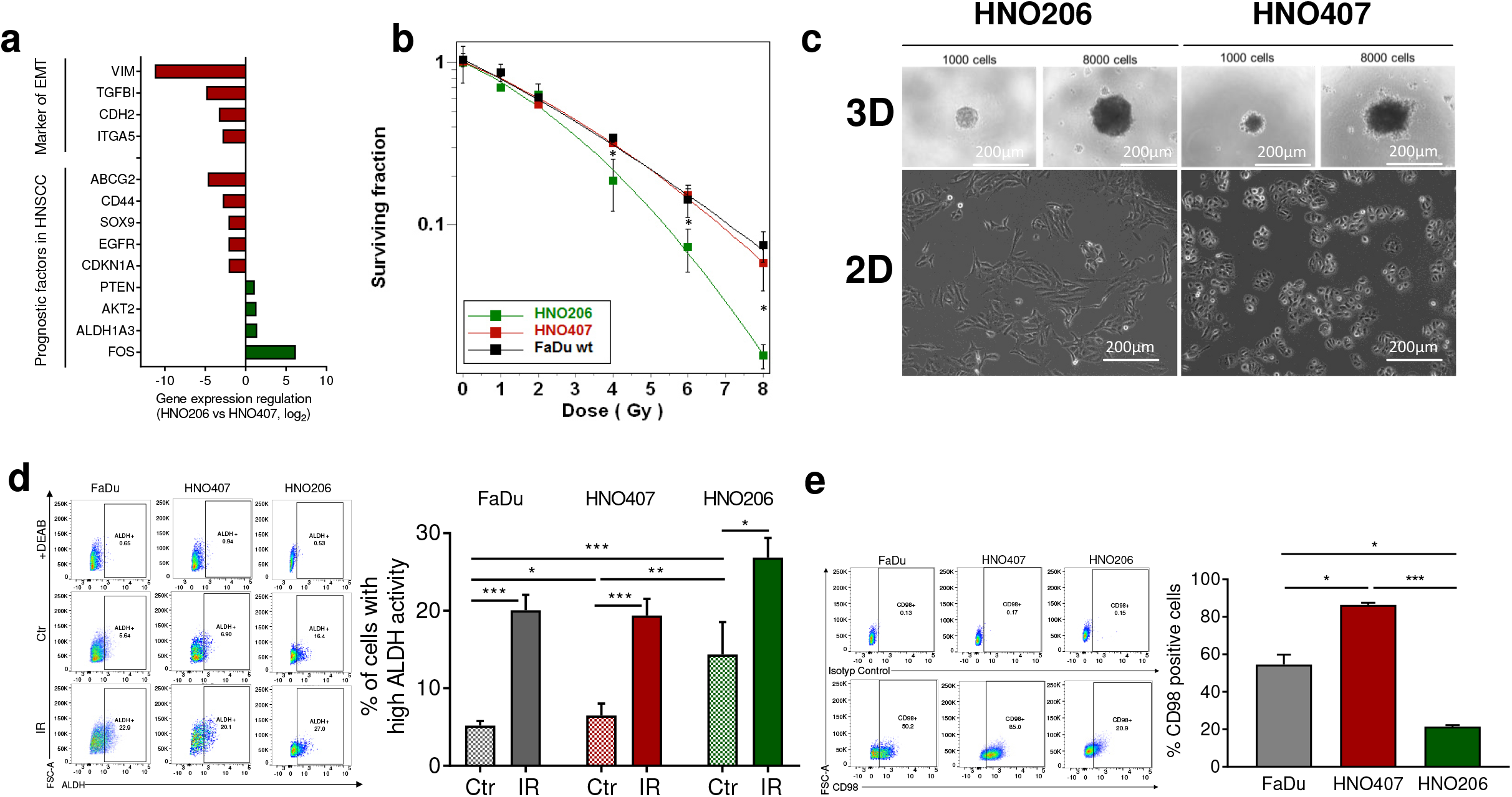
The expression of HNSCC specific biomarkers correlates to the radiation sensitivity of the patient-derived HNSCC lines analyzed. A Expression regulation of genes involved in EMT and genes suggested as prognostic factors for overall survival of patients with HNSCC. Unpaired t-test was performed by comparing HNO407 to HNO206 (unpaired t-test, p<0.01). B Radiobiological clonogenic survival assay comparing the relative radiosensitivity of FaDu (black), HNO407 (red) and HNO206 (green) (*, p<0.05). C Morphological appearance of patient-derived HNO407 and HNO206 cells grown in a 2D monolayer and in a 3D spheroid culture (representative phase-contrast microscopic images, scale bar = 200μm). D Representative flow cytometry plots of ALDH activity in FaDu, HNO407 and HNO206. ALDH activity was assessed using the ALDUFLUOR assay under non-irradiated conditions (Ctr) and after fractionated irradiation (IR) of 5 x 4Gy, measured 48h after the last irradiation fraction. Data shown is one representative replicate of three independent replicates. Gates for calculating high ALDH activity were placed according to the control sample treated with the ALDH inhibitor diethylaminobenzaldehyde (DEAB). Bar plot shows the flow cytometric analysis of ALDH activity under non-irradiated conditions (Ctr) and after fractionated IR in all cell lines (**, p<0.001; ***, p<0.0001, error bars = SD). E Representative flow cytometry plots showing the proportion of CD98 positive cells in FaDu, HNO407 and HNO206 under non-irradiated conditions (Ctr). Gates for CD98 positive cells were placed based on the mouse IgG isotype control sample. Data shown is one representative sample of three independent samples. Bar plots summarize the flow cytometric analysis of the expression of CD98 under non-irradiated conditions in all cell lines (*, p<0.05; **, p<0.001; ***, p<0.0001, error bars = SD).

The radiation sensitivity of the selected HNSCC tumor cell lines was assessed in clonogenic survival assays using increasing doses of X-ray. In repetitive experiments it was observed that the radiation sensitivity of the patient-derived HNO407 line shows no significant differences compared to the FaDu line, whereas the HNO206 line showed a significant (p<0.05) increase in radiation sensitivity of 1.8-fold (**Fig. 2B**). The HNO206 and HNO407 lines also showed morphological differences in 2D and 3D cell culture models (**Fig. 2C**). HNO206 cells display a more elongated cell morphology compared to HNO407 cells which have a polygonal morphology in 2D cultures (**Fig. 2C**). In 3D spheroid cultures of HNO407 exhibited an irregular contour with loosely attached cells at the border of the spheroids. In comparison, HNO206 cells formed densely packed spheroids with strongly adhering cells. The frequency of two biomarkers, Aldehyde dehydrogenase (ALDH)^24,25^ and CD98^26,27^ that have been previously reported to be associated with increased radiation resistance in HNSCC cell lines and poor prognosis in patients with HNSCC were evaluated by flow cytometry. Differences in the proportion of cells with high ALDH activity was measured across the HNSCC cell lines. The highest frequency of tumor cells with a high ALDH activity was measured in HNO206 cells of which 14% showed high capacity to oxidize aldehydes. In the HNO407 line this population was decreased to 7% and in the FaDu line to 5% (**Fig. 2D**). This correlated to the ALDH1A3 expression levels analyzed by RNA-seq analyses (**Supplementary Data 1B**). Flow cytometry was in addition performed to determine the frequency of cells expressing the cell surface marker CD98. In the HNO407 patient-derived line 86% of the tumor cells were CD98 positive. In the FaDu line 55% were CD98 positive followed by the radiation sensitive patient-derived line HNO206 in which 22% were CD98 positive (**Fig. 2E**). The proportion of CD98 positive cells correlates with the radiosensitivity of the cell lines (R^2^=0.87, **Supplementary Fig. 1B**). In previous studies it has been reported that cells with high ALDH activity increase upon exposure to fractionated radiation^25,28^. We evaluated these observations in the reference FaDu line and compared this with the patient-derived HNSCC lines after 5 x 4Gy fractionated radiation as described previously^28^. In the FaDu line we could confirm a 3.9-fold increase in cells with high ALDH activity after radiation. Cells with high ALDH activity in the HNO206 line were increased 1.8-fold and in the HNO407 line 3.0-fold (**Fig. 2D, Supplementary Fig. 1C**). Although the frequency of ALDH positive cells under nonirradiated conditions did not correlate to the degree of radiation sensitivity the induction of ALDH activity after fractionated radiation therapy was the strongest in the less sensitive FaDu and HNO407 line (**Supplementary Fig. 1D**). The increase of CD98 positive cells in response to fractionated radiation was in contrast anti-correlating in respect to radiation sensitivity. In the high intensity CD98 positive fraction which in the HNO206 line resembled a small subpopulation below 1% we observed the strongest increase in CD98 positive cells (**Supplementary Fig. 1E, F**).

### High resolution lentiviral cellular barcoding of HNSCC lines enables longitudinal tracing of clonal dynamics in response to irradiation

To explore the dynamics of individual tumor cell clones and their progeny in response to IR the three HNSCC cell lines were clonally marked with the previously described high-complexity lentiviral ClonTracer barcode library^18^. The library consists of 10^7^ semi-random 30 nucleotide-long DNA sequences encoded in lentiviral vectors that serve as unique barcodes for the genomic marking of individual cells within a heterogeneous cell population. Upon transduction with the ClonTracer barcode library and integration of the lentiviral DNA into the cell genome, tumor cells are labelled with uniquely identifiable DNA barcodes. The barcodes are passed onto daughter cells upon cellular division enabling high-resolution quantitative cell fate tracking and deconvolution of clonal population dynamics by multiplex PCR-amplification of barcodes and subsequent next generation sequencing (NGS). To achieve an accurate cell marking, the FaDu and the two patient-derived cell lines HNO206 and HNO407 were transduced with the ClonTracer library with a low multiplicity of infection (MOI 0.1) ensuring that each tumor cell is clonally marked by only one lentiviral vector and its corresponding barcode. Antibiotic selection was performed to select tumor cells that have been efficiently transduced and contained barcodes in their genome. For subsequent experiments barcode-labelled tumor cells were expanded for 14 days to achieve an approximately 20-fold representation of each barcode. The total number of barcoded cells was evaluated after NGS by bioinformatic processing as previously described^18^.

The number of detected barcodes is directly proportional to the number of individual clones, which were marked by the lentiviral vector. We could confirm the clonal marking of 1.2×10^5^, 1.9×10^5^ and 1.6×10^5^ FaDu, HNO206 and HNO407 cells, respectively (**Supplementary Fig. 2A**). The quantification of sequencing reads for each barcode provides information on the clonal composition and abundance of each clone. Based on the quantification of all barcoded clones in the three tested cell lines we could confirm that after 14 days of cell expansion no major differences in the clonal abundance of individual clones was detectable. The sequencing reads of unique barcodes in each tumor cell line were equally distributed in all barcoded tumor cell lines verifying that no significant clonal skewing took place in the expansion phase and before the initiation of the experiments (**Supplementary Fig. 2B**). Furthermore, we evaluated the resolution of our detection method by comparing the clonal composition of two technical replicates for the HNSCC patient-derived lines after NGS. Therefore, clonal barcodes were amplified and sequenced from a total of 5×10^6^ clonally marked cells in two independent technical replicates showing an close to 80% overlap of detected clones (**Supplementary Fig. 2C**).

### Fractionated exposure to ionizing radiation results in clonal selection and reduction in clonal complexity

To monitor clonal dynamics of HNSCC cell lines and the contribution of individual tumor cell clones to clonogenic survival in response to IR, a repetitive clonogenic survival assay with the ClonTracer barcoded tumor cell lines was performed. Barcoded cells were exposed to fractionated IR with 5 fractions of 4Gy and were seeded for a clonogenic assay 48h after the last fraction (**Fig. 3A**). To avoid stochastic loss of barcodes and to ensure representation of all barcodes in each replicate, clonogenic assays were performed in replicates with 4.4×10^6^, 4.6×10^6^ and 1.9×10^6^ cells, the total of cells which survived exposure to fractionated radiation in the FaDu, HNO407 and HNO206 line, respectively. The number of cells that survived fractionated IR confirmed the low radiosensitivity of the HNO206 line compared to HNO407 and FaDu in clonogenic assays (**Supplementary Fig. 3A, B**). In parallel, clonal expansion of tumor cells in a clonogenic assay was monitored under non-irradiated conditions.

**Figure 3 -.**
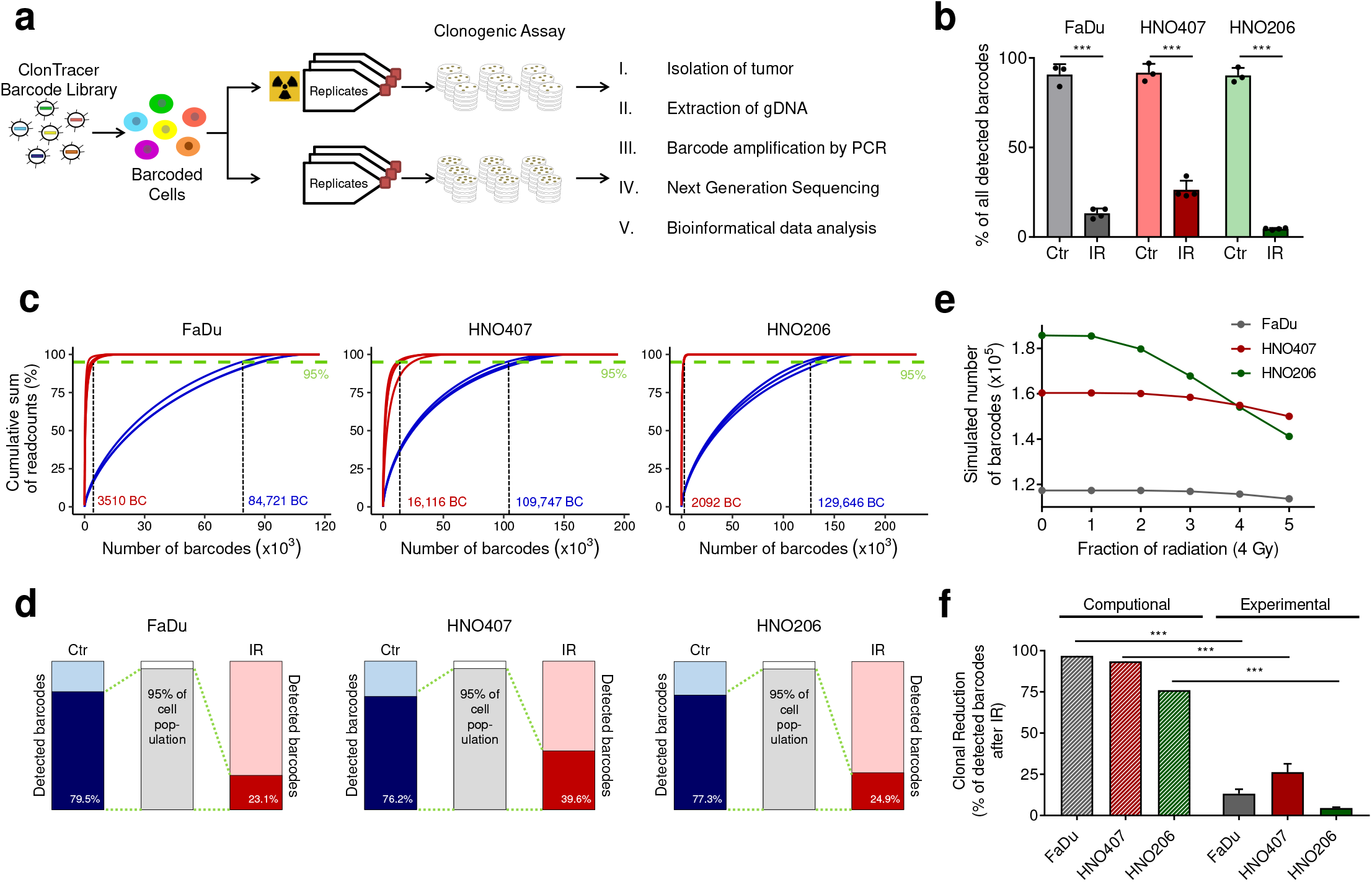
Fractionated irradiation promotes reduction in clonal complexity. A Experimental overview. Upon transduction of the HNSCC cell lines with the lentiviral ClonTracer barcode library, cells were expanded and fractionated irradiated with 5 fractions of 4Gy or maintained under normal non-irradiated conditions. 48 h after last irradiation cells were seeded in multiple 150 mm2 dishes in low density for colony formation. Colonies were harvested as a clonal pool and barcode sequences were PCR-amplified from genomic DNA (gDNA) and subjected to NGS. NGS data was processed by in-house software for subsequent analysis of clonal dynamics. B Bar plots showing the average proportion of detected barcodes after colony formation in nonirradiated (Ctr, n=3) and irradiated (IR, n=4) samples in FaDu (gray), HNO407 (red) and HNO206 (green). Percentages are calculated relative to the number of barcodes detected in the initial starting population of the respective cell line (***, p<0.0001, error bars = SD). C Cumulative sum of barcode sequencing reads of non-irradiated (blue) and irradiated (red) samples in FaDu, HNO407 and HNO206. The average number of barcodes that constitute 95% of total barcode sequencing reads under non-irradiated and irradiated conditions is highlighted. D Bars demonstrate the average proportion of detected barcodes in control (Ctr, blue) and irradiated replicates (IR, red) that contribute to 95% of the total barcode sequencing reads. E Numerical modeling of the number of detected barcodes after each fraction of radiation in FaDu, HNO407 and HNO206 assuming that all cell clones have the same radiation survival probability and the same capability to repopulate after each treatment cycle. F Bar plots showing the average model-predicted proportion of detected barcodes after 5 fractions of photon radiation (4Gy) and experimentally detected barcodes in FaDu (gray), HNO407 (red) and HNO206 (green), (n =4; ***, p<0.0001, error bars = SD).

In the absence of irradiation, in principle every clonally marked tumor cell had clonogenic potential determined after repeated seeding. In average 90.2 - 91.9% of clonally marked cells in all three analyzed tumor cell lines had the capacity to seed and expand in colonies. The capacity of tumor cells to efficiently seed and to clonally expand was generally decreased after exposure to fractionated IR (**Fig. 3B, Supplementary Fig. 3C**). In the patient-derived HNO206 line, which showed the highest degree of radiation sensitivity, in average 4.5% of the initially clonally marked clones were able to seed and clonally expand after exposure to fractionated IR, which is equivalent to a 22-fold reduction in clonal complexity. In contrast, clonal reduction in FaDu and HNO407, which showed an increased radiation resistance, was less intensive as compared to the HNO206 line. In the FaDu line in average 13.2% of the initial clonally marked cells seeded and formed colonies resembling an 8-fold reduction in clonal complexity. In the HNO407 line, clonally marked tumor cells showed the highest capacity to seed and clonally expand with in average 26.3% of the initially marked cells correlating to a 4-fold reduction in clonal complexity after exposure to fractionated radiation (**Fig. 3B, Supplementary Fig. 3C**).

The reduction in clonal complexity was in addition verified by the cumulative sum of barcode sequencing reads that constitute 95% of total barcoded population (**Fig. 3C**). In the nonirradiated FaDu line 95% of all reads was reached with ~84,721 unique barcodes and was reduced to ~3510 unique barcodes after fractionated IR. In the radiation sensitive HNO206 line this reduction was even more severe dropping from ~129,646 barcodes under non-irradiated conditions to ~2092 barcodes after exposure to IR. The HNO407 line, in contrast, revealed the lowest degree of clonal reduction ranging from ~109,747 barcodes under non-irradiated conditions to ~16,116 barcodes after irradiation (**Fig. 3C**). Accordingly, in the non-irradiated FaDu line 79.5% of barcodes contributed to 95% of total barcode sequencing reads compared to 23.1% after IR (**Fig. 3D**). In non-irradiated HNO407 and HNO206 line, 76.2% and 77.3% of detected barcodes constituted 95% of the total barcode sequencing reads, which dropped to 39.6% and 24.9% in irradiated HNO407 and HNO206, respectively (**Fig. 3D**).

To analyze whether the observed clonal reduction in response to IR is lower or higher than expected, we applied a numerical simulation to calculate the expected clonal sizes after fractionated radiotherapy. The simulation provides expected clonal sizes based on experimental data such as barcode representation, total number of clonally marked cells, survival fraction at 4Gy and number of radiation fractions for each tumor cell line. Further, the simulation assumes that all cell clones have the same radiation survival probability and the same capability to repopulate after each treatment cycle. Accordingly, in independent simulations the number of remaining barcodes decreased after each irradiation fraction in all cell lines, however, to a significantly lesser extent than observed experimentally (**Fig. 3E**). Clonal reduction after the last fraction was calculated and the total barcode number was reduced by 3.1% in the FaDu line. In the HNO407 line the simulation calculated a reduction by 6.5% and in HNO206, which showed the highest sensitivity to radiation, a reduction of 24.0% was calculated by numerical simulation (**Fig. 3F**). This contradicts what we observed after fractionated radiation experimentally, which resulted in the clonal reduction of 86.8%, 73.7% and 95.5% of the initial barcodes in the FaDu, HNO407 and HNO206 lines, respectively. Thus, the experimentally measured clonal reduction significantly exceeds what would be expected assuming equal radiosensitivity and repopulation capacity across all barcoded cells. This intensity of clonal reduction correlates to the radiosensitivity and to the proportion of CD98 positive cells (R^2^>0.76, **Supplementary Fig. 3D, E**).

### Clonogenicity after exposure to radiation is characterized by intensive clonal skewing of individual tumor cells

The sequencing reads of each barcode is directly proportional to the clonal population size of the individual tumor cell clone (**Supplementary Fig. 4**). This enables a dissection of clonal abundancy and a comparison of clonal expansion rates for each cell and their progeny under non-irradiated and irradiated conditions. We reasoned that with this analysis we can first determine the capability of each clone to survive radiation and second evaluate the clonogenicity of each clone after exposure to IR (**Supplementary Fig. 4**). The frequency distribution of barcodes under non-irradiated conditions showed a Gaussian distribution, which flattened and shifted in response to fractionated IR increasing the pool of low and high abundant tumor cell clones (**Fig. 4A**). To determine the extent of clonal skewing and distortion, the sequencing reads were normalized to the initial starting barcode population and log_2_-transformed to obtain values for increased and decreased clonal abundance in comparison to the initial starting population. Clonal skewing as a consequence of fractionated IR was observed in all cell lines, while the control populations showed a symmetrical distribution (**Supplementary Fig. 5A**). The proportion of clones with high and low abundancy was quantified with fixed thresholds based on 4-fold (log_2_) enrichment or depletion in clonal abundancy compared to the initial starting population. Accordingly, in the clonally marked FaDu line that was exposed to IR ~12.5% were identified as low and ~12.2% as high abundant clones. These values were significantly (p<0.05) lower under non-irradiated conditions where low abundant clones were detected with a frequency of ~1.1% and high abundant clones at a frequency of ~2.0%. Similar clonal shifts were detectable in the two patient-derived HNSCC lines. In the HNO407 line, 7.7% low abundant and 8.5% high abundant clones were detected after exposure to IR while under non-irradiated conditions these values were 2.4% for low abundant and 1.8% for high abundant clones (p<0.05). In the irradiated HNO206 line, 8.8% of the clonally marked cells showed low abundance and 24.7% were highly abundant. In this line under non-irradiated conditions 3.2% of the clones were low and 2.5% high abundant (p<0.05) (**Fig. 4B, Supplementary Fig. 5B**). These characteristics in clonal dynamics of irradiated tumor cells in clonogenic survival indicate an extensive clonal distortion and skewing upon exposure to IR. This is defined by a small fraction of tumor clones, which preferentially survive irradiation while maintaining their clonogenic potential to dominantly expand after exposure to IR.

**Figure 4 -.**
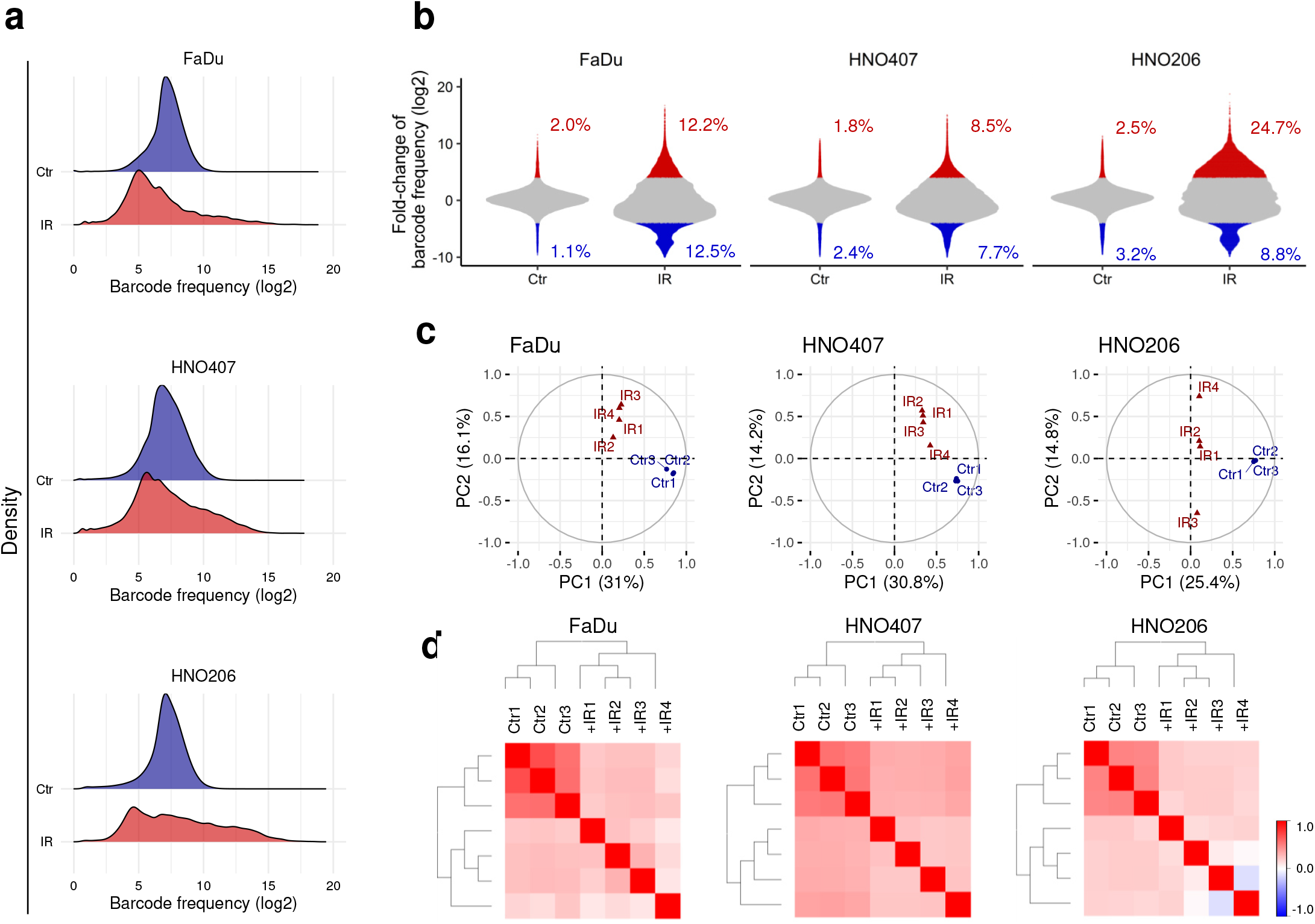
Survival after exposure to radiation is characterized by intensive clonal skewing of individual tumor cells. A Barcode frequency distribution of the non-irradiated (blue) and irradiated population (red) represented by log_2_ transformed sequencing reads. The frequency distributions show one representative distribution of 3 control samples and 4 irradiated samples. B Violin plots illustrate the density distribution of all detected barcodes in a representative replicate and indicate the average proportion of 4-fold enriched (red) and 4-fold depleted (blue) (log_2_) barcodes in non-irradiated (Ctr) samples and irradiated (IR) samples after normalization to the initial starting population in FaDu, HNO407 and HNO206. C Principal component analysis (PCA) on the patterns of clonal composition resulted in 2 distinct groups of non-irradiated (Ctr) and irradiated (IR) samples. Percentage variation explained per coordinate of Principal Component 1 (PC1) and Principal Component 2 (PC2). D Hierarchical clustering of technical replicates of barcoded FaDu, HNO407 and HNO206 distinguishes between non-irradiated and irradiated samples (Pearson Correlation, Manhattan distance).

To detect patterns of clonal composition and to evaluate clonal correlations among nonirradiated and irradiated replicates we applied a principal component analyses (PCA). The PCA separated the non-irradiated and irradiated replicates of each cell line into two distinct groups (**Fig. 4C**). The grouping was confirmed by hierarchical clustering on the basis of a Pearson correlation (Manhattan distance), which resulted in two distinct cluster branches in all three cell lines consisting of non-irradiated control and irradiated replicates (**Fig. 4D**). Furthermore, Pearson correlation coefficients were pairwise calculated to quantify the correlation of the clonal compositions between non-irradiated and irradiated replicates. For non-irradiated FaDu replicates, an average Pearson correlation coefficient of 0.61 was determined, which was significantly reduced to 0.18 (p<0.001) after irradiation (**Supplementary Fig. 5C**). For the HNO407 line the average Pearson coefficient was 0.54 for non-irradiated replicates reaching 0.23 (p<0.001) after exposure to IR. For HNO206 cells, the Pearson coefficient of non-irradiated replicates was 0.48 decreasing to 0.08 (p<0.001) (**Supplementary Fig. 5C**). In comparison, the Pearson correlation coefficient among irradiated replicates in the radiosensitive cell line HNO206 decreased 6.0-fold after IR, whereas in FaDu and HNO407 the decrease was 3.4- and 2.3-fold, respectively. The similarity of clonal composition of irradiated replicates correlates with the cellular radiosensitivity and CD98 expression of the different cell lines (**Supplementary Fig. 5D, E**, R^2^>0.96). The less radiosensitive HNO407 line, which has the highest proportion of CD98 expressing cells, shows the highest similarity among the clones which survived radiation, whereas the radiosensitive HNO206 line shows the weakest similarity among clones which have survived radiation.

### Clonal dynamics of radiation survival unveil stochastic and deterministic parameters of radiation survival in HNSCC cell lines

Up to the present it is poorly understood whether survival of cancer cells after exposure to radiation is driven by clones, which have intrinsic molecular characteristics enabling them to preferentially survive exposure to radiation in a deterministic manner. In contrast, radiation itself may have the potential to change the degree of functional heterogeneity, which could mechanistically influence the probability of cancer cells to survive radiation in a rather stochastic manner. We reasoned that by clonally marking tumor cells and analyzing the survival probability and clonogenicity of every clone and its progeny in response to radiation, deterministic and stochastic parameters of radiation survival and clonogenicity could be determined within different cancer cell populations (**Supplementary Fig. 4**). We hypothesized that detecting the same clonal barcodes in two or more independent irradiated experimental replicates, the associated tumor clones could potentially harbor intrinsic mechanisms that enable them to preferentially survive the exposure to IR. This group of tumor cells we defined as the deterministic population in relation to radiation survival and clonogenicity. In contrast, the survival and expansion of tumor cells after exposure to IR could also be driven by different clones in a stochastic manner. In such a scenario, clones that survive and clonally expand would be expected to be different between experimental replicates. To quantify the population within tumor cells lines with stochastic survival probabilities we determined the number of clonal barcodes that are unique for each replicate. In opposite, to quantify the population with deterministic survival probabilities the number of clonal barcodes that are detected in at least two replicates was evaluated.

While under non-irradiated conditions in all HNSCC lines the majority of clonogenic clones was shared between replicates this changed upon exposure to fractionated irradiation dramatically. In the FaDu line 36.0% of the individually marked clones represented the deterministic tumor cell population, which preferentially survive exposure to radiation whereas 64.0% of the clones survived in a stochastic manner (**Fig. 5A, Supplementary Fig. 6A**). In the HNO407 line the highest frequency of a deterministic tumor cell population was measured with 45.6% of the clonal barcodes found to survive radiation in more than one replicate. In the radiation sensitive HNO206 line the lowest frequency of a deterministic tumor cell population was observed with 17.8% of the clonal barcodes found in more than one replicate (**Fig. 5A, Supplementary Fig. 6A**). The observed frequency of shared barcodes is significant (p<0.001) based on the Fisher’s exact test for all HNSCC lines analyzed and not a result of random sampling.

**Figure 5 -.**
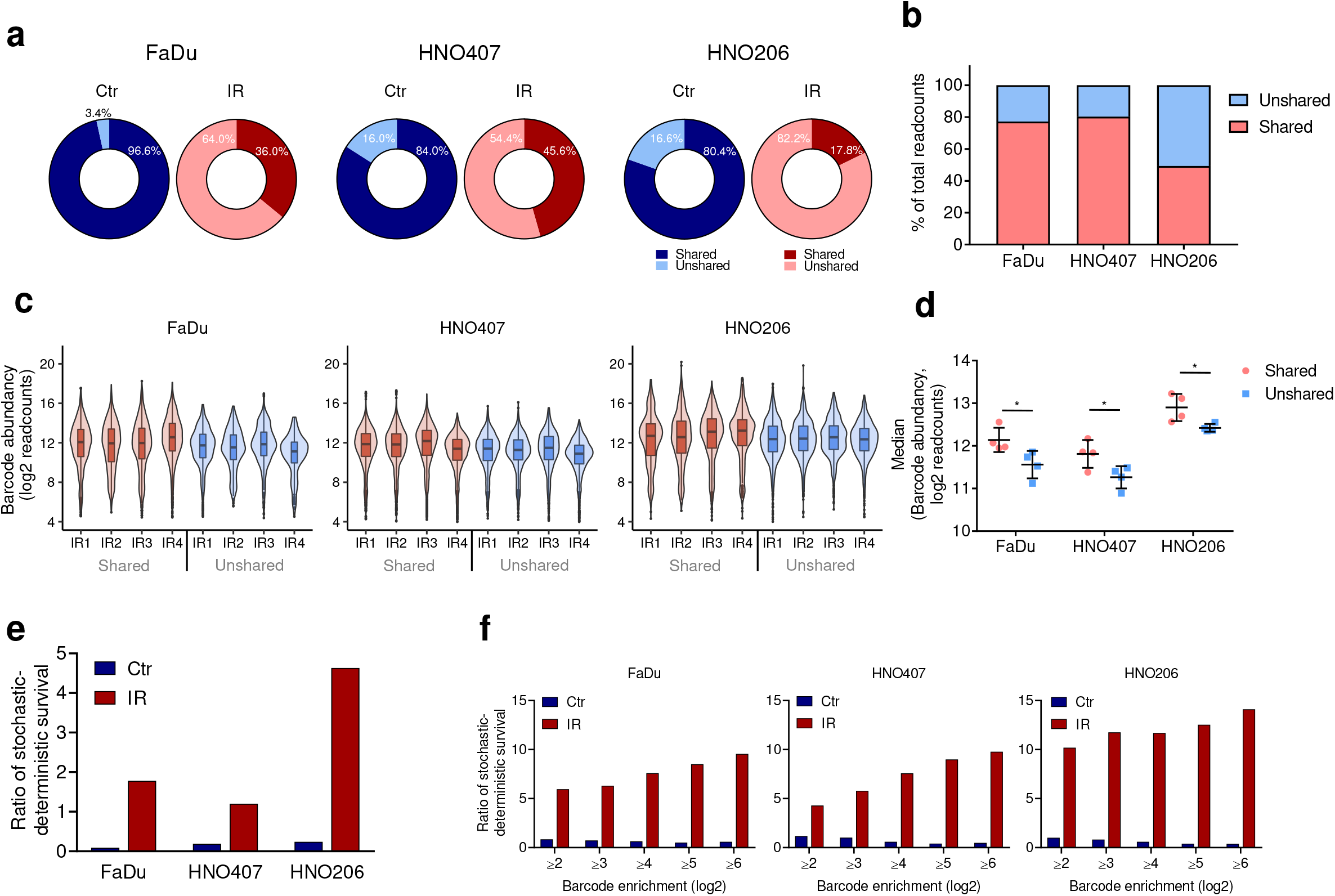
Determining stochastic and deterministic survival parameters in response to irradiation. A Pie charts display the percentage of unshared barcodes and barcodes shared in at least two replicates in non-irradiated control (Ctr, blue) and irradiated samples (IR, red) in FaDu, HNO407 and HNO206. All barcodes that passed bioinformatic processing were considered. B Stacked bar chart represents the proportion of the total barcode sequencing counts represented by unshared or shared barcodes in FaDu, HNO407 and HNO206. C Violin- and boxplots represent ≥4-fold enriched barcodes (log_2_) detected in only one replicate (unshared barcodes) or in at least two replicates (shared barcodes). D Average median of readcounts (log_2_) of ≥4-fold enriched barcodes (log_2_) detected in deterministic cell population (shared barcodes, red) and in stochastic cell population (unshared barcodes, blue), (unpaired t-test, *, p<0.05). E Stochastic-deterministic survival ratios for all detected barcodes F Stochastic-deterministic survival ratios for for barcodes with different thresholds of abundancy for non-irradiated (Ctr, blue) and irradiated (IR, red) barcoded FaDu, HNO407 and HNO206 cells. The stochastic-deterministic survival ratio was calculated by the division of the proportion of unshared barcodes by the proportion of shared barcodes.

### The stochastic-deterministic survival ratio in patient-derived HNSCC lines provides insights in the frequency of intrinsically radiation resistant tumor cell clones

The clonogenic capacity for each tumor cell clone can be quantified by the sequencing count of each barcode (**Supplementary Fig. 4**). This provides quantitative knowledge on the overall clonal size and the intensity of the clonal expansion within the stochastic and deterministic cell population. By quantifying the sequence counts of both populations following exposure to fractionated irradiation we can first evaluate the survival probability and second define the clonogenic capacity of both populations by introducing thresholds for the relative clonal abundance of each clone. Based on this we can show that survival, the pure presence of clones following exposure to radiation, is preferentially driven by the deterministic tumor cell population although this population has less absolute clone numbers (**Supplementary Fig. 4**). In the FaDu and HNO407 line 77.2% and 80.3% of the barcode readcounts originated from the deterministic population. In the radiosensitive HNO206 line in contrast, the capacity of the deterministic population to survival was significantly lower at 49.3% of the total sequencing counts (**Fig. 5B**). Following selection of clones which survive and seed after exposure the clonogenic capacity of tumor cell clones was measured for each population based on the sum of sequencing reads of each clone. The degree of clonogenicity was determined by increasing sequencing read thresholds (**Supplementary Fig. 6B**). With increasing thresholds the contribution of the deterministic population to clone sizes and clonogenicity decreases in all three HNSCC lines. This indicates that in opposite to survival probabilities the clonogenic capacity is not predominantly driven by the deterministic population but mainly by the stochastic population (**Supplementary Fig. 6B**). Despite the fact that the proportion of deterministic clones decreases after IR in all cell lines, the relative clonogenic capacity is increased in the deterministic (shared) compared to the stochastic (unshared) tumor cell population (**Fig. 5C, D**, p<0.05). This indicates that the clonogenic capacity within the deterministic population is preserved contributing to a significant clonal proportion of the survival fraction in clonogenic assays.

We aimed to evaluate whether stochastic or deterministic probabilities of tumor cell clones to survive radiation could be used to predict the frequency of intrinsically radiation resistant clones in patient-derived tumor cell populations. Therefore, we determined a stochastic-deterministic survival ratio based on clone numbers identified after exposure to irradiation. The proportion of non-shared clones, which were only detected in a single replicate, was divided by the proportion of shared clones, which were found in 2 or more replicates representing the deterministic population that commonly survives exposure to radiation (**Supplementary Fig. 7A**). The ratio provides a measure to define whether radiation survival and clonogenicity in a tumor cell population follows a stochastic (ratio >1) or a deterministic route (ratio <1). In all cell lines the stochastic-deterministic ratio for the non-irradiated population was smaller than 1 (FaDu: 0.04, HNO407: 0.19, HNO206: 0.24, **Fig. 5E**). Following exposure to IR the ratio shifted to values were the number of clones, which had survived radiation in only one replicate succeeded the number of clones, which had commonly survived radiation. The stochastic-deterministic survival ratio showed a linear correlation to the radiation sensitivity and proportion of CD98 positive cells of the lines analyzed (**Supplementary Fig. 7B**, R^2^=0.99, **Supplementary Fig. 7C**, R^2^=0.89). This shows in support of our hypothesis that radioresistant cell lines have a higher proportion of pre-existing clones that harbor intrinsic molecular mechanisms to survive radiation. The HNO206 line, which showed the highest radiation sensitivity also had the highest stochastic-deterministic ratio of 4.62 followed by the FaDu line with 1.78 and 1.19 for the HNO407 line (**Fig. 5E**). The preferential stochastic clonal selection after exposure to fractionated IR was also visible while analyzing clone numbers, which were enriched in their abundance above a particular threshold (**Fig. 5F**). Interestingly, rare tumor cells that commonly survived exposure to IR in all replicates was detected at a frequency of 1.4%, 3.3% and 0.64% for FaDu, HNO407 and HNO206, respectively (Fishers Exact test, p<0.001). The observed frequency of detecting such clones was not the result of random sampling since compared to the statistically expected values it was 1.4–4.4-fold higher than to the frequency of commonly detected clones in non-irradiated control samples (**Supplementary Fig. 7D**). This indicates that the frequency of clones which survive radiation in a deterministic manner is not the result of a random selection providing evidence for the existence of a rare subpopulation with an intrinsically radiation resistant phenotype commonly surviving radiation. From insights in these clonal dynamics we conclude that survival probabilities in contrast to clonogenic capacities have to be seen as two different stages within clonogenic assays. Survival is mainly driven by a tumor cell population which has a higher probability to resist radiation and is deterministically selected. The clonogenic capacity in contrast is a result of clonal competition and selection of the fittest clones which is preferentially driven in a stochastic manner (**Fig. 6**).

**Figure 6 -.**
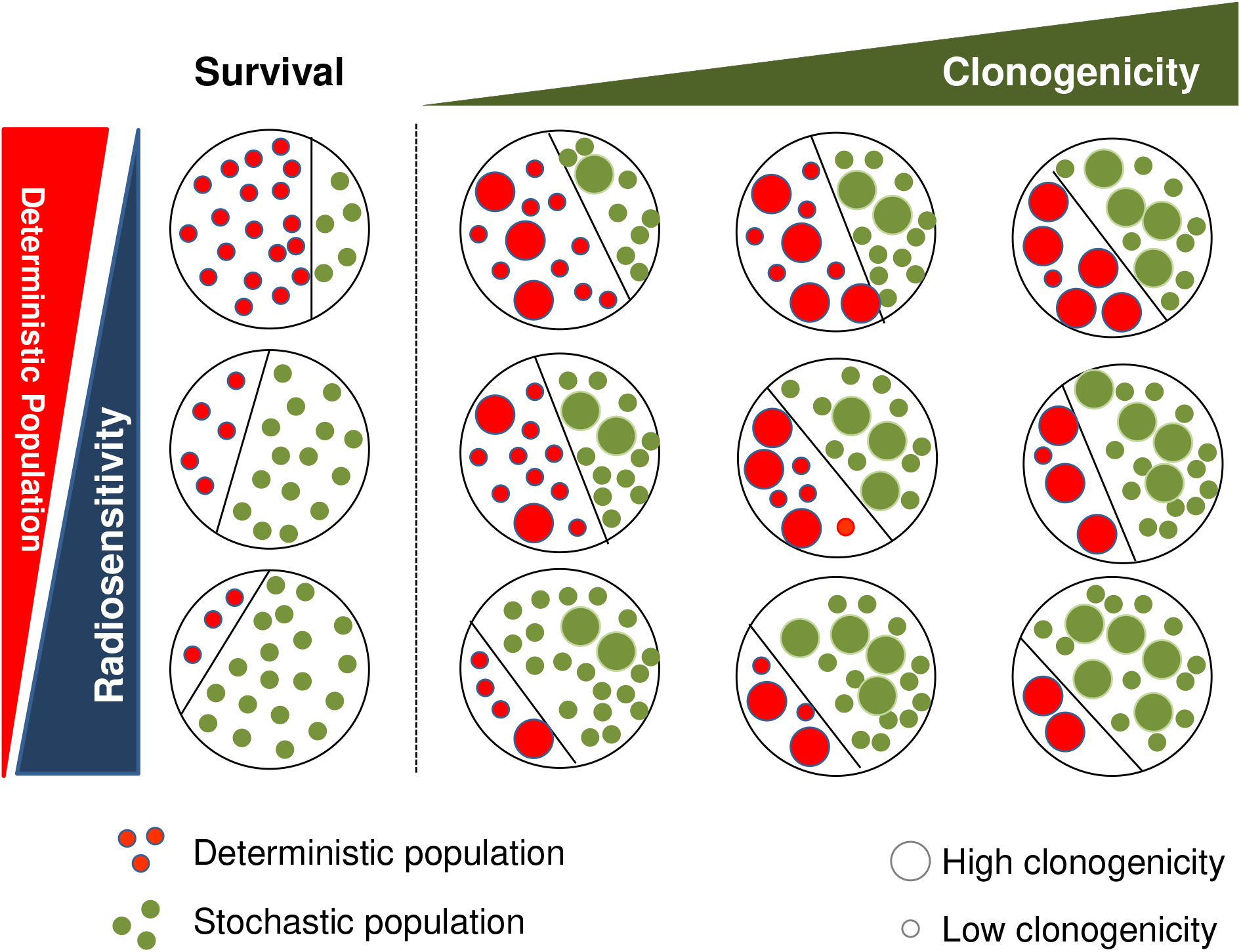
Graphical abstract of clonal dynamics in response to radiation. The frequency and contribution of deterministic and stochastic tumor cell population to survival and clonogenicity in relation to the decrease of radiation sensitivity is illustrated.

## Discussion

The clonogenic survival assay is an essential method to determine the sensitivity of cells to ionizing radiation or anti-cancer drugs^2,29^. The determination of the surviving fraction assumes a homogenous radiosensitivity across individual tumor cell clones of the cell population thereby masking the impact of functional heterogeneity, which potentially exists in regards to the radiosensitivity of individual tumor cells. Radiation survival and clonogenic potential may have strong clone-to-clone variations, which are currently overlooked in standard clonogenic assays. In this context it is not known whether the surviving fraction can be characterized by tumor cell clones, which either have intrinsic or acquired capacities to survive radiation.

To address these questions, we have made use of a lentiviral based cellular barcoding strategy in HNSCC patient-derived cell lines and introduced a new qualitative value to the readout of radiobiological clonogenic survival assays. This value consists of deciphering and quantifying clonal dynamics of radiation survival and clonogenicity in parallel in single cell resolution using NGS. With cellular barcoding of the FaDu and two HNSCC patient-derived cell lines using the previously published ClonTracer barcode library^18^ the clonal fate of more than 1×10^5^ cells per in three different patient-derived HNSCC lines were quantified in multiple replicates of clonogenic assays in response to fractionated IR. This enabled a comprehensive analysis in the clonal dynamics of tumor cells in response to IR.

The capacity of cells to survive radiation and to retain clonogenic potential decreases with increasing doses of IR. The degree of clonal reduction correlated with the radiosensitivity of the respective HNSCC line analyzed and was significantly higher than numerical modeling suggested if survival probabilities within a tumor cell population would be uniform. The experimentally derived values on clonal reduction therefore support the fact that profound clone-to-clone differences in the sensitivity to radiation exist within tumor cell lines. The majority of tumors cells fail to survive exposure to radiation. The small fraction of cells which have survived exposure to fractionated irradiation thereby indeed show a higher survival probability and have been clonally selected based on their low radiation sensitivity. It can be excluded that the probability of an inhomogeneous cell irradiation, which is due to fluctuations in both, the number of tracks passing the target cell and the energy deposition per track^30^ influences this observation, since fractionated irradiation was performed increasing the chance that each cell gets equivalent doses across the experiment.

Clonal reduction was concomitant with an extensive clonal skewing within the surviving fraction. This indicates that following survival to radiation a competition in clonogenic capacities between the surviving clones exists, which is most probably influenced by differences in relative fitness of each clone after exposure to IR. Clonal skewing was defined by a dominant subpopulation, which not only preferentially survived exposure to radiation but showed a high clonogenic capacity. In contrast the surviving fraction also contained tumor cell clones which showed weaker clonogenic potential. The ability of individual tumor cells to survive exposure to IR and to retain clonogenic potential can therefore be considered as independent characteristics with strong clone-to-clone variations as some surviving clones hold intensive clonogenicity whereas others stay in a rather dormant, non-proliferative condition.

We hypothesized that if this clonogenic survival after exposure to radiation is a stochastic process determined by different clones which have acquired the ability to survive radiation, the clonal barcode architecture after radiation will be different across replicates. By contrast, if clonogenicity after exposure to radiation is driven deterministically by clones harboring intrinsic molecular mechanisms promoting radiation survival these clones should be detected in more than one replicate. In non-irradiated samples the majority of barcodes were detected in more than one replicate. Clonal dynamics of tumor cells in response to irradiation however indicate that radiation survival and clonogenicity can be seen as two different selection steps. In the first step the probability of tumor cells to survive exposure to irradiation is dominated by the clonal selection of a smaller deterministic population. In the second step the clonogenic capacity is a result of clonal competition with cells which have survived radiation stochastically and which numerically form the larger tumor cell population within the survival fraction. By quantifying the frequency of the stochastic and deterministic subpopulations in clonogenic assays a stochastic-deterministic survival ratio was generated which enabled to compare the clonal composition between different HNSCC lines. The frequency of the deterministic subpopulation contributing to the survival fraction in clonogenic assays correlates to the degree of radiation sensitivity and the expression level of HNSCC biomarker CD98. The less radiosensitive HNO407 line has the lowest and the radiosensitive HNO206 line the highest ratio. This observation in turn highlights that in the radiosensitive line the stochastic radiation survival is more pronounced, while the less radiosensitive line HNO407 is composed of a higher proportion of cells with intrinsic molecular features which commonly survive IR in a deterministic manner. This is supported by the fact that a frequency of rare tumor cells that always survived exposure to IR was determined which was significantly higher than expected by random sampling increasing from 0.64 to 3.3% linearly to the degree of radiation sensitivity. It can be speculated that this tumor cell population have a truly intrinsically radiation resistant phenotype which warrants further molecular investigation of this population using single cell gene expression analyses in a similar experimental set up.

We show that high resolution cellular barcoding for longitudinal tracing of tumor cells in response to irradiation and the implementation of a stochastic-deterministic survival ratio provide a new qualitative value to oversee the impact of functional heterogeneity in clonogenic assays. The here defined parameters for distinguishing tumor cell populations, which survive radiation in a stochastic and deterministic manner and enable cross comparison of patient-derived tumor lines in respect to radiation resistance on the clonal level. Deciphering the clonal identity of tumor cells which contribute to survival and clonogenicity in response to radiation will help to devise future strategies to characterize possible differences between acquired and intrinsic molecular mechanisms that promote radiation resistance.

## Author Contributions

A.N. conceived the project, supervised and designed the experiments. AW designed and performed the experiments. C.S developed and executed bioinformatical processing tools and computer simulation and advised on statistics and data analyses. C.P provided valuable discussions. I.K provided resources and valuable discussion. C.H.M kindly provided the patient-derived cell lines and provided valuable discussions. A.A. and J.D. provided resources, valuable discussion and conceptual advice. A.W and A.N analyzed the data and wrote the manuscript.

## Acknowledgements

For technical assistance, we are grateful to Chante Mayer and Lea Günther. For next-generation sequencing, we would like to thank the core facility of the DKFZ. Funding was provided to A.N, I.K and C.P by the joint funding program of the National Center for Radiation Oncology (NCRO) and Heidelberg Institute of Radiation Oncology (HIRO) as well as intramural funds of the national center for tumor diseases (NCT) personalized radiation oncology (PRO) program.

## Material and Methods

### Cell lines and cell culture

The HNSCC cancer cell line FaDu was purchased from the American Type Cell Collection (ATCC) and cultured in Minimum Essential Medium (MEM, Gibco™/Life Technologies) with 10% Fetal Bovine Serum (FBS, Sigma-Aldrich) and 1% Penicillin and Streptomycin (10,000U/mL / 10mg/mL; Gibco™/Life Technologies). HNO206 is a patient-derived tumor cell line from a primary HNSCC tumor. HNO407 is a patient-derived cell line derived from a lymph node metastasis. Patient-derived cell lines were kindly provided by Prof. Dr. Chistel Herold Mende, University Clinic Heidelberg. Cell lines were genotyped using microsatellite polymorphism analysis and regularly confirmed to be free of mycoplasma. Patient-derived cell lines were cultured in Dulbeccos’s Modified Eagle Medium (DMEM, Biochrom) with 10% FBS and 1% Penicillin and Streptomycin. Cells were cultured at 37°C in a humidified environment and 5% CO2 atmosphere. Cultures were regularly tested to be free from mycoplasma using Venor^®^GeM Classic Kit (Minerva Biolabs). Single-cell suspensions for experiments were obtained from exponentially growing cultures by mild enzymatic detachment using a 0.05% trypsin / 0.02% EDTA solution in phosphate buffered saline (PBS, PAN Biotech). 3D spheroids were generated applying the Liquid Overlay procedure in agarose-coated 96 well plates^31^. Shortly, 96-well plates were coated with 50μl of 1.5% agarose-solution (complete medium, 1.5% agarose). 2000 cells per well in 200μl complete medium were seeded and cultured for 4 days for spheroid formation.

### ClonTracer library lentivirus production

ClonTracer pooled plasmid library was purchased from Addgene^18^, Addgene #69830). ClonTracer lentivirus was produced by lipofection using Lipofectamine LTX with Plus Reagent (Invitrogen) in HEK-293T cells. 2.5×10^6^ HEK-293T cells were seeded in in 10cm cell culture dishes. For each 10cm dish, 2.4μg ClonTracer plasmid DNA, 2.4μg psPAX2 (encoding HIV-1 Gag, Pol, Tat and Rev proteins) and 0.6μg pMD2.G (encoding VSV G envelope protein) was diluted in 2000μl Opti-MEM^®^ Reduced Serum medium (Gibco™/ Life Technologies). 5.4μl PLUS™ Reagent was added directly to the diluted DNA and incubated for 5min at RT. 16.2μl Lipofectamine™ LTX Reagent was added to the diluted DNA and incubated for 30min at RT. DNA-Lipofectamine™ LTX complexes were added dropwise to the cells. 16h post-transfection medium was replaced by 6ml complete medium (10% FBS, 1% Penicillin and Streptomycin). 40h after transfection using a syringe with 0.45μm filters. Virus suspension was aliquoted and stored at −80°C.

### Titration and barcoding of cancer cells

For lentiviral barcoding we use a low multiplicity of infection to ensure that cells are labelled with one single barcode and to reduce the chances of any single cell being marked with more than one barcode. In order to find optimal virus dilution for each cell line to achieve a MOI of 0.1, the virus was titrated on each cell line. Therefore, 0.5×10^6^ cells were seeded into a 6-well plate with complete medium (10% FBS, 1% Penicillin and Streptomycin). After cell attachment, different dilutions of virus suspension were added (1:10, 1:100, 1:1000, 1:10000 virus dilution). 24h posttransduction medium was replaced by 2ml complete medium. 48h after transduction cells were detached and splitted as replicates into two new 6-well plates. Medium containing 2μg/ml puromycin was added to one replicate. 72h after puromycin addition cells were harvest and counted using a haemocytometer. Percent transduction was determined by dividing cell count from the replicate with puromycin by cell count from the replicate without puromycin multiplied by 100. The virus amount was determined that achieved approximately 10% cell survival (MOI 0.1) and used for clonal barcoding of cells. Consequently, the virus titer of the virus suspension was determined. Virus titer was calculated by using the following formula.

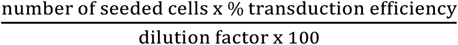

Cells were barcoded by lentiviral infections with the ClonTracer lentivirus using a target MOI of 0.1 that corresponds to 10% of infectivity after puromycin selection. Therefore, totally 12×10^6^ cells were plated into 12 wells of a 12-well plate (1×10^6^ cells/well) and after cell attachment, the respective volume of lentivirus suspension was added. 16h post transduction cells were trypsinized, combined and replated into 2 T175 cell culture flasks. 72h later medium was replaced by 2μg/ml puromycin containing medium for 72h to select for successfully transduced cells. Infected cell populations were expanded in culture for 2 weeks to obtain sufficient number of cells to set up replicate experiments.

### Clonogenic survival assays

Determination of radiosensitivity in clonogenic survival assays were performed in 6-well plates. A low density cell number (50-800 cells/well) was seeded that increased with escalation of radiation dose. Irradiation (MultiRad irradiation device by Faxitron, 200 kV, 17.8 mA, 0.5 mm Copper filter) was performed 16h after cell seeding. After 12-14 days depending on cell line doubling time colonies were washed and fixed to the plate surface using fixation solution (75% methanol, 25% acetic acid). After 10min of incubation at RT fixed colonies were washed twice with dH_2_O. Staining solution (0.1% Chrystal violet in dH_2_O) was added and plates were incubated for 10min at RT to stain colonies. Plates were washed twice with dH_2_O and dried over night for plate scanning and subsequent analysis. The images were analyzed with ImageJ and clonogenic graphs were created and analyzed using in-house software CS-Cal (implemented by Dr. C Schwager, http://angiogenesis.dkfz.de/oncoexpress/software/cs-cal). Colony counts were normalized to non-irradiated samples for visualization of cellular survival curves and for determination of surviving fractions. Each experiment was carried out in triplicates/group and was repeated three times.

To monitor clonal dynamics after fractionated irradiation in clonogenic assays, 5×10^6^ barcoded cells were fractionated irradiated (MultiRad irradiation device by Faxitron) on 5 consecutive days receiving 4Gy (200 kV, 17.8 mA, 0.5 mm Copper filter) per day. 48h after receiving the last fraction, cells were seeded in 150mm^2^ cell culture plates with a density of 0.8×10^6^ cells/plate. Non-irradiated control cells were seeded with a density of 0.1×10^6^ cells/plate. Cell culture plates were incubated for a period of 7-14 days, depending on doubling time. Medium was changed every second day. Once colonies were formed, cells were washed with PBS and trypsinized to harvest cells. Cells were washed with PBS and 5×10^6^ cells were snap-frozen for genomic DNA extraction and RNA isolation.

### Flow cytometric analyses

Flow cytometry analysis was applied for the quantification of cell surface markers and determination of ALDH activity. Antibody concentrations were determined in titration experiments to achieve maximum cell staining with minimum antibody concentration. Respective immunoglobulin (IgG) isotype-matched controls were used as negative control to measure nonspecific background staining. For labelling the cell surface marker CD98, cells were mechanically harvested and the single-cell suspension was washed twice with staining buffer (PBS, 2% FBS, 0.1% sodium azide). 0.25×10^6^ cells were resuspended in 200μl of staining buffer containing anti-CD98 antibody (CD98-Alexa680, Santa Cruz Biotechnology, Inc., sc-59145) or respective isotype control (mouse IgG-Alexa680, Santa Cruz Biotechnology, Inc., sc-516621) and incubated for 45min at 4°C. Propidium iodide (PI; 2μg/ml) was added for dead-cell discrimination and incubated for 5min. Cells were washed twice and resuspended with staining buffer for subsequent flow cytometric analysis on BD FACSCanto II. Analysis of the proportion of positively stained cell populations was performed using FlowJo™ software (Version 10.3). To determine intracellular ALDH activity the Aldefluor^®^ Kit (Stem Cell Technologies) was used according to manufacturer’s instructions. The optimal amount of Aldefluor^®^ substrate was determined in titration experiments to achieve maximum cell staining with minimum Aldefluor^®^ substrate concentration. Shortly, cells were harvested and 0.5×10^6^ single cells were resuspended in 1ml Aldefluor^®^ buffer. 5μl of Aldefluor^®^ substrate was added and mixed briefly. 500μl of the cell suspension was immediately transferred to another tube supplemented with ALDH inhibitor diethylaminobenzaldehyde (DEAB), which is used as negative control for background fluorescence. Cells were incubated 45min at 37°C. During incubation the Aldefluor^®^ substrate (BODIPY-aminoacetaldehyde, BAAA) is converted by the ALDH enzyme into a green fluorescent product (BODIPY-aminoacate, BAA) that is retained in the cell and can be detected in FITC channel. After incubation cells were washed with Aldefluor^®^ buffer containing TOTO™-3 Iodide (Invitrogen™) to counterstain and exclude dead cells in subsequent flow cytometric analysis. After 5min of incubation cells were washed and resupsended in 500μl Aldefluor^®^ buffer for flow cytometric analysis on BD FACSCanto II. Analysis of the proportion with high ALDH activity was performed using FlowJo™ software (Version 10.3). To determine the proportion of CD98 high expressing cells and the ALDH activity after fractionated IR, 2.5×10^6^ cells were seeded in T75 cell culture flasks. After 16h cells were fractionated irradiated (MultiRad irradiation device by Faxitron) on 5 consecutive days receiving 4Gy (200 kV, 17.8 mA, 0.5 mm Copper filter) per day. 48h after receiving the last fraction, cells were harvested and stained as described above for subsequent analysis.

### Simultaneous purification of DNA and RNA

Genomic DNA (gDNA) and total RNA were extracted using Quiagen AllPrep DNA/RNA Mini Kit according to manufacturer’s protocol. In brief, 5×10^6^ cells were disrupted by adding 600 μl buffer RLT Plus and subsequent vortexing. For homogenization, the lysate was passed 5 times through a 20-gauge needle fitted to an RNase-free syringe. The lysate was transferred to an AllPrep DNA spin column and centrifuged. The gDNA bound column was stored at 4°C for later DNA purification. 600μl 70% ethanol was added to the RNA-containing flow-through and transferred to an RNeasy spin column. After washing, total RNA was eluted from the column membrane by adding 50μl nuclease-free water. Genomic DNA bounded membrane was washed and gDNA was eluted in 50μl nuclease-free water. DNA and RNA concentrations were determined using Nanodrop spectrophotometer, gDNA was stored at −20°C, and RNA was stored at −80°C.

### Barcode amplification

Barcodes were amplified by PCR using Phusion Flash high fidelity PCR Mastermix (Thermo Fisher Scientific) with primers containing adaptors for Illumina sequencing and 10-bp-long index sequences to allow sample multiplexing (Vector forward primer: “actgactgcagtctgagtctgacag”, Vector reverse primer: “ctagcatagagtgcgtagctctgct”). The barcodes were amplified in a nested PCR to increase specificity of the PCR reaction. The sampling of sufficient template coverage was ensured by 8 parallel PCR reactions for each sample in the first and second PCR run. 1.5μg of gDNA was used as a template in the first PCR run of the nested PCR for each PCR reaction. PCR reactions were pooled and 5μl of the pool was used as a template for each reaction of the second PCR run. PCR products were size selected and cleaned up using AMPure XP PCR Purification solution (Beckman Coulter™). After purification, PCR products were dissolved in 100μl nuclease-free water and DNA concentration was determined using Nanodrop spectrophotometer. PCR products were stored at −20°C.

### Next-generation sequencing

500ng of PCR product per sample was pooled and PCR product pool was quantified on Qubit 4 Fluoremeter using Qubit dsDNA BR Assay Kit (Thermo Fisher Scientific) and finally diluted to a 10nM solution for sequencing on Illumina HiSeq2500 (FaDu) or Illumina HiSeq4000 (HNO407 and HNO206) sequencer. Sequencing was performed at the DKFZ Genomics and Proteomics Core Facility provding results of the barcode-sequencing in FastQ files.

### Barcode processing and analysis

To prepare sequencing readcount tables for barcode analysis, FastQ files were processed using the in-house developed software SeedScan (Dr. Christian Schwager, https://angiogenesis.dkfz.de/oncoexpress/software/seedscan/index.htm) according to guidelines described before^18^. In brief, ClonTracer barcodes from multiplexed samples were identified based on matched forward/reverse multiplex index primer sequence allowing 1bp error and assigned to the specific sample library. ClonTracer barcodes were excised and filtered according to 1) barcodes showed the expected barcode nucleotide pattern of 15x WS (Weak=nucleotide pair with 2 hydrogen bonds (A,T), Strong=nucleotide pair with 3 hydrogen bonds (G,C)). 2) Barcodes matched the expected index primer sequence (“agcagag”), which is comprised of the last bases of the ClonTracer 3’ vector directly after the 15x WS barcode sequence. 3) Barcode is detected with at least 100 readcounts in two samples and 4) barcodes are detected in the initially barcoded cell population. To compare barcode numbers, barcode frequencies and barcode composition across samples, sequencing readcounts of each sample were normalized to the totally measured sequencing reads (**Supplementary Data 2**). To further analyse foldchanges in barcode frequency the sequencing readcounts were normalized to the sequencing readcounts of the initially detected barcode population and log_2_ transformed. Pearson Correlation analysis was used to study similarities of barcode composition across replicates and subsequent data clustering was performed by Manhattan distance using the in-house software SUMO (Dr. Christian Schwager, https://angiogenesis.dkfz.de/oncoexpress/software/sumo/). Principal component analysis to study similarity of replicates was performed with the factoMineR and factoextra R package in R statistical software.

### Significance of barcode overlaps by Fisher’s Exact test

In order to examine the intersection between detected barcodes of independent replicates as well as their statistical significant applying Fisher’s Exact^32^ test we used the Venn analysis tool from the in-house developed software SUMO. Venn analysis tool provides a number of intersection between two lists of barcodes that is expected when the selection occurs randomly from the same base population (Null hypothesis; p-value ~1). In case the experimentally observed intersection between two replicates is much differently than the calculated expected intersection, the intersection cannot be explained by random sampling and is statistical significant (p≪1). To analyze the significance of barcode intersection observed for all possible combinations of shared barcode intersections between the non-irradiated control and irradiated samples, p-values based on Fishers exact test were determined. Shared barcodes were selected as those, which were present in at least two replicates. Further, to compare the intersection of 3 samples in the control and IR group in consideration of the different number of barcodes that were detected in the group, we calculated the ratio of observed and expected barcode overlap based on the Fishers Exact test. The expected overlap of 3 samples was determined for control (n=3) and for every possible combination of the three irradiated replicates (n=4). The experimentally observed barcode intersection was normalized to the mean of the calculated expected barcode intersection providing a ratio of observed to experimentally expected barcode overlap (**Supplementary Fig. 7D**).

### RNA sequencing, processing and analysis

RNA sequencing was performed on FaDu, HNO407 and HNO206 (3 biological replicates). Total RNA was isolated from snap-frozen cells by using Quiagen AllPrep DNA/RNA Mini Kit (see simultaneous purification of DNA and RNA) and quantified using Agilent RNA 6000 Nano Kit (Agilent Technologies) on the Agilent 2100 Bioanalyzer. 1.2μg high purity total RNA (defined as having an RNA integrity number (RIN) greater than 8.0) was used as input for RNA seq library construction. RNA seq library construction and paired-end sequencing reading length 100bp on HigSeq4000 was conducted at the DKFZ Genomics and Proteomics Core Facility. FastQ files from sequencing where mapped to the human genome and transcripts were counted. TopHat v2.1.1 was used for mapping applying default settings^33–37^. Cufflinks v2.2.1 was used for counting of transcripts applying default settings^38^. Counts are reported as Fragments per Kilobase transcript length per Millions reads analysed (FPKM).Tools were downloaded and installed with conda/miniconda 4.7.12 (https://docs.conda.io/projects/conda/en/latest/index.html) on Ubuntu 16.04.6 LTS Linux system (https://ubuntu.com/). Reference genome Homo_sapiens.GRCh38.83 as well as respective GTF file (description of transcripts) were downloaded from NCBI’s ftp server (https://ftp.ncbi.nlm.nih.gov/). Finally, cufflinks’ gene counts of individual samples were combined into an expression matrix for further statistical analysis. Statistical analysis was carried out using the in-house software SUMO (Dr. Christian Schwager, https://angiogenesis.dkfz.de/oncoexpress/software/sumo/). Therefore, RNA transcripts with an average count >10 were considered. Normalization and log_2_ transformation was performed for the analysis of differentially expressed genes. To identify genes differentially expressed in two samples, unpaired t-test was carried out. To identify genes differentially expressed in more than two sample groups, analysis of variance (ANOVA) was performed. Although RNA-Seq expression counts may be not Gaussian distributed, Gaussian based tests (t-test, ANVOA) should sufficiently rank genes as more/less significantly differential expressed to filter significant genes for further system biology analysis, whereas the resulting p-value may be not exact. Pathways affected by differentially expressed genes were identified using STRING (https://string-db.org/, Reactom data base)^39^ and Reactome^40^ (https://reactome.org/). Gene set enrichment analysis^41,42^ (GSEA, https://www.gsea-msigdb.org/gsea/index.jsp) was performed to identify differentially regulated pathways using the Hallmark gene set^43^. Statistically significant gene sets are defined with nominal p-value<0.05 and FDR q-value<0.05.

### Simulation

In order to recapitulate whether the clonal reduction after fractionated irradiation can be observed, when assuming that each barcoded cell has the same radiosensitivity and growth ability, we established a numerical simulation. The numerical simulation is fed with experimentally derived data such as the surviving fraction at 4Gy (FaDu: 0.298, HNO407: 0.312, HNO206: 0.214), the number of cells in the experiment (5×10^6^) and the total number of detected barcodes (FaDu: 117303, HNO407: 160392, HNO206: 185798). After each fraction of radiation, the simulation calculates the number of barcodes that remains based on the surviving fraction. Until the next fraction of radiation, remaining barcoded cells are able to divide until reaching maximum cell number (5×10^6^). This simulation is repeated for five fractions of radiation providing a number of remaining barcodes after each fraction.

### Statistics

Data are represented as means and standard deviations (SD). Statistical significance was determined two-tailed Student’s t-test in GraphPad Prism (Version 7, GraphPad, San Diego, CA, USA). The asterisks represent significantly different values (*, p<0.05; **, p<0.001; ***, p<0.0001). Data represent average values of at least three independent replicates. In ClonTracer clonogenic survival experiments, we had three independent replicates for the nonirradiated control and four independent replicates for fractionated irradiation. In experiments with more than two sample groups, analysis of variance (ANOVA) was performed.

## Supplementary information

### Supplementary Figure legends

**Supplementary Figure 1.**
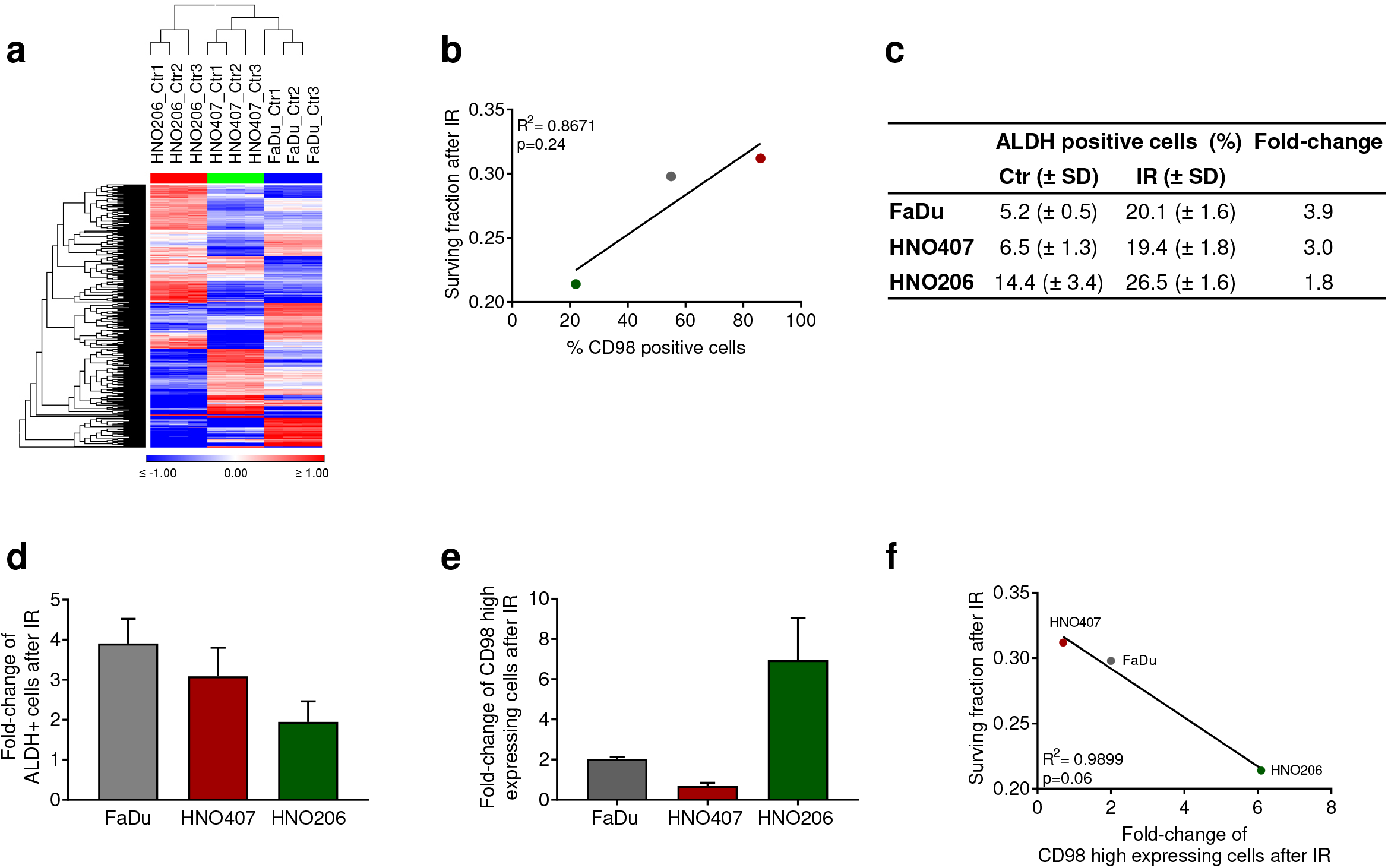
(**a**) Heatmap showing differentially regulated genes in FaDu, HNO407 and HNO206 (1-factor ANOVA, p<0.00005). Hierarchical clustering (Manhattan distance) revealed that HNO206 clustered separately from FaDu and HNO407. (**b**) Linear correlation analysis of the surviving fraction of FaDu, HNO407 and HNO206 at 4Gy with the proportion of CD98 positive cells (R^2^=0.8671, p=0.24). (**c**) Table shows the average proportion of ALDH positive cells in FaDu, HNO407 and HNO206 in non-irradiated cells (Ctr) and after fractionated IR of 5 x 4Gy measured 48h after the last fraction of radiation (Mean ± SD). (**d**) Average fold-change of ALDH+ cells after 5 fractions of 4Gy (unpaired t-test, *, p<0.05; n.s. not significant, error bars=SD). (**e**) Average fold-change of CD98 high expressing cells after 5 fractions of 4Gy (unpaired t-test, *, p<0.05; **, p<0.001; ***, p<0.0001, error bars = SD). (**f**) Linear correlation analysis of the fold-change in CD98 high expressing cells and the surviving fraction after 4Gy (R^2^=0.9996, p=0.01).

**Supplementary Figure 2.**
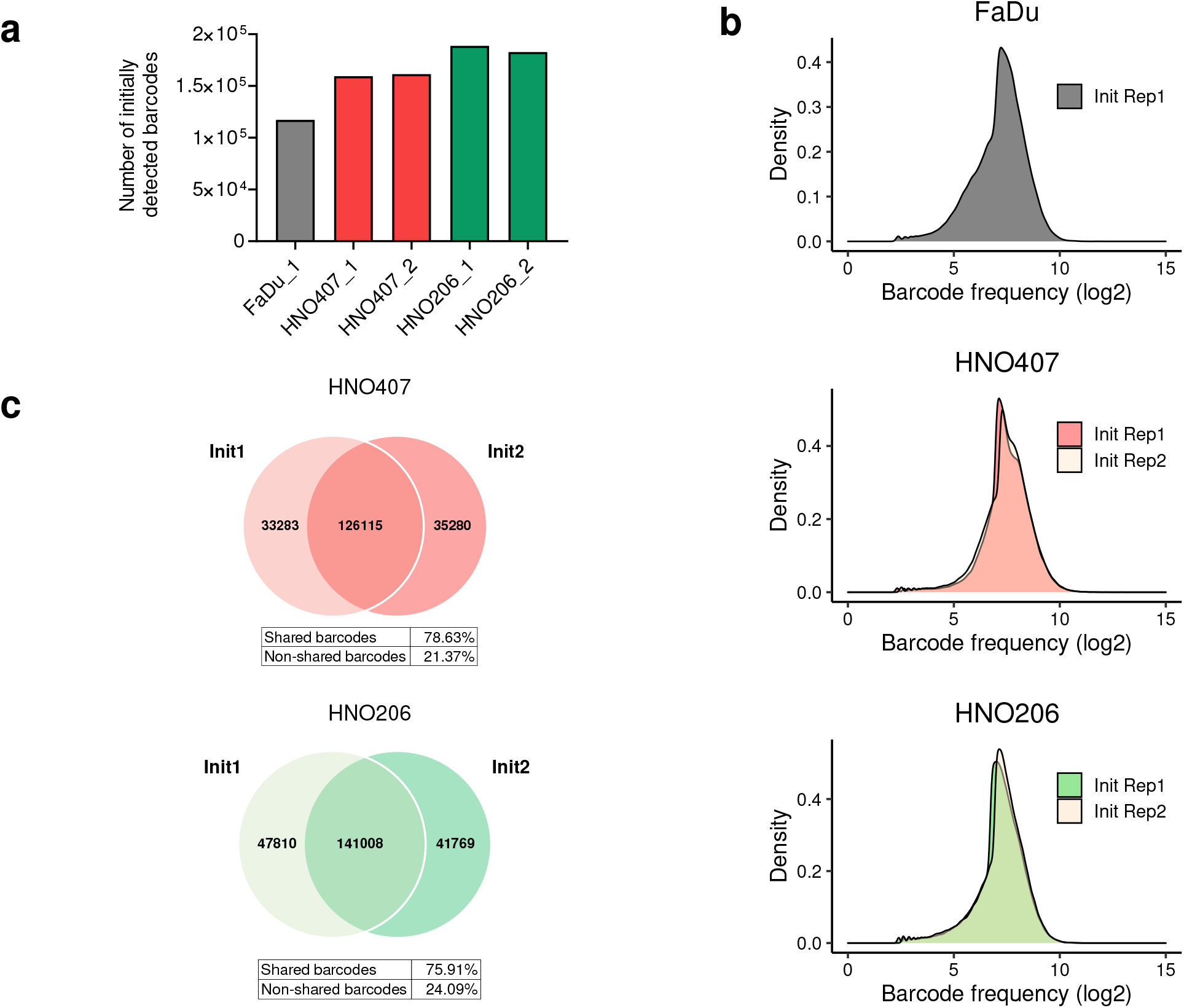
(**a**) Bar plots showing the number of detected barcodes in the initial starting population of FaDu, HNO407 and HNO206 after transduction with the ClonTracer lentiviral barcode library, antibiotic selection, expansion and bioinformatic processing. One technical replicate of initial starting population was processed for the FaDu line and two technical replicates for the HNO407 and HNO206 line. (**b**) Distribution of the barcode sequencing reads (log_2_) of the initial starting population in FaDu and of two technical replicates of the initial starting populations of HNO407 and HNO206. (**c**) Venn diagram displaying the number of overlapping barcodes found in each initial starting population of HNO407 and HNO206. All detected barcodes after bioinformatic processing were included. Tables show percentages of shared and non-shared barcodes among the two processed initial starting populations.

**Supplementary Figure 3.**
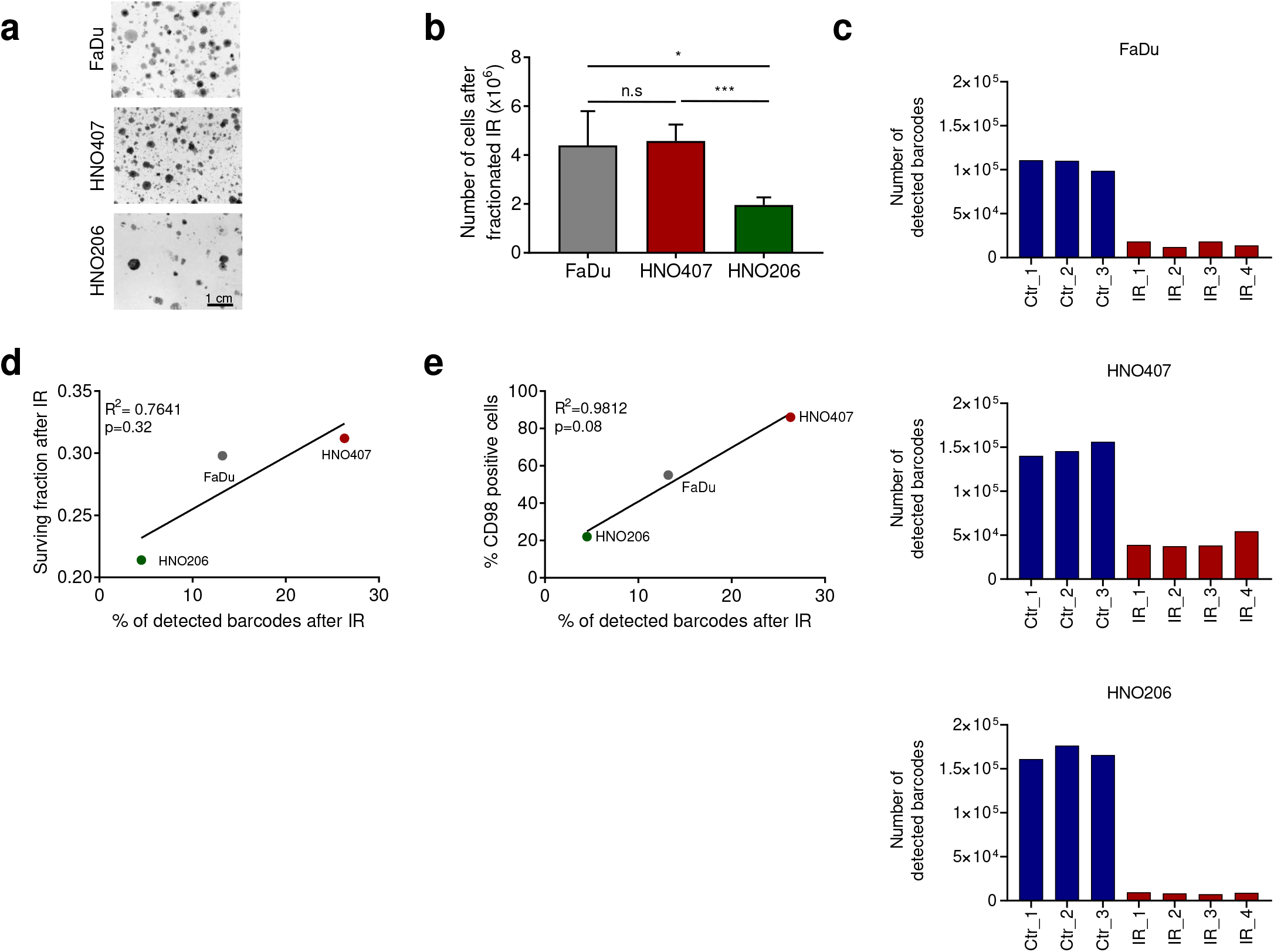
(**a**) Representative images of colonies formed after 5 fractions of photon radiation with 4Gy in FaDu, HNO407 and HNO206. (**b**) Number of cells that survived fractionated IR (5×4Gy) and were seeded for clonogenic assays (unpaired t-test, *, p<0.05; ***, p<0.0001; n.s., not significant, error bars = SD). (**c**) Bar plots showing the number of detected barcodes after colony formation in non-irradiated control samples (Ctr, blue) and samples that were exposed to fractionated photon radiation (5×4Gy, IR, red) in FaDu, HNO407 and HNO206. (**d+e**) Linear correlation analysis of the proportion of detected barcodes after fractionated IR of FaDu, HNO407 and HNO206 and (**d**) the surviving fraction at 4Gy (R^2^=0.7641, p=0.32) and (**e**) the proportion of CD98 positive cells (R^2^=0.9812, p=0.08).

**Supplementary Figure 4.**
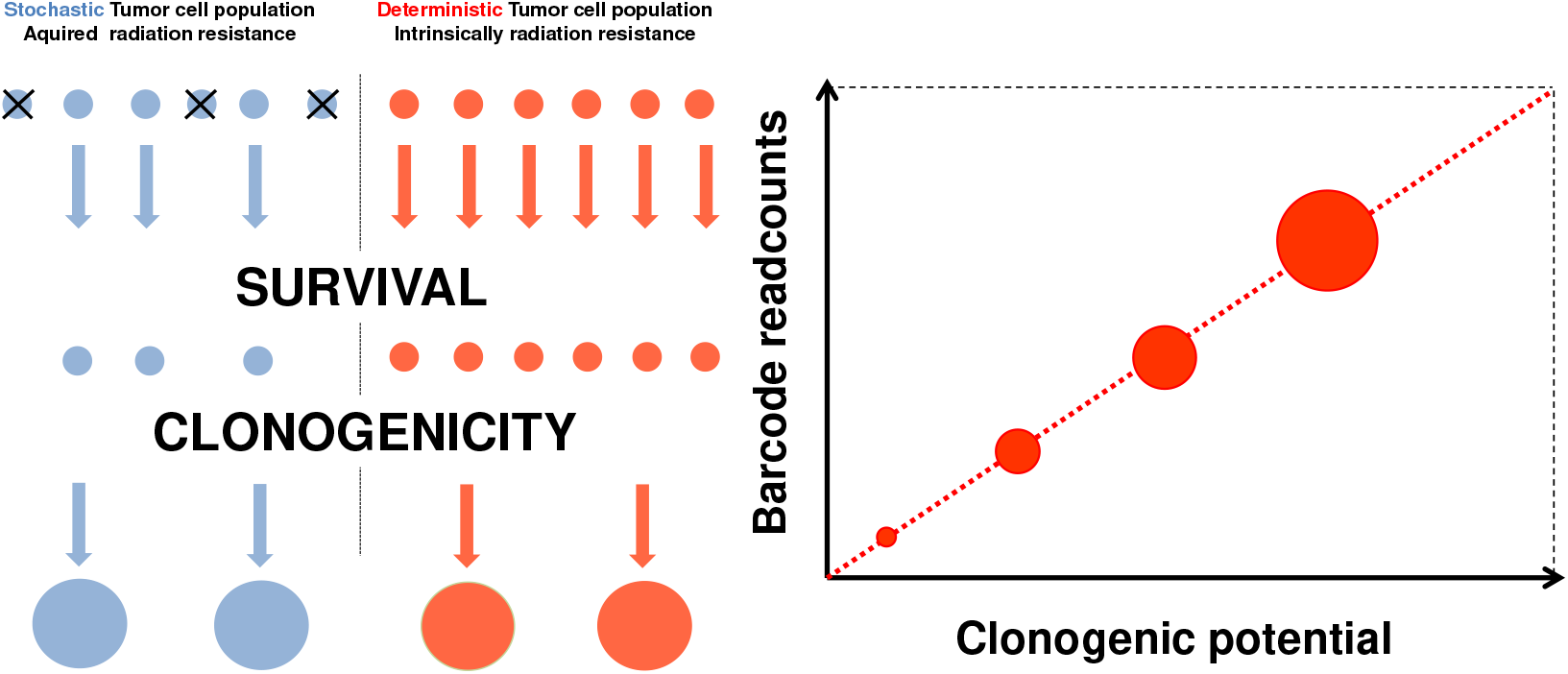
Graphical model for the deterministic and stochastic tumor cell populations that survive exposure to radiation and preserve their clonogenic potential. The clonogenic potential for each tumor cell clone is directly proportional to the sequencing counts of the respective barcode providing quantitative knowledge on the overall clonal size and the intensity of the clonal expansion within the stochastic and deterministic cell population.

**Supplementary Figure 5.**
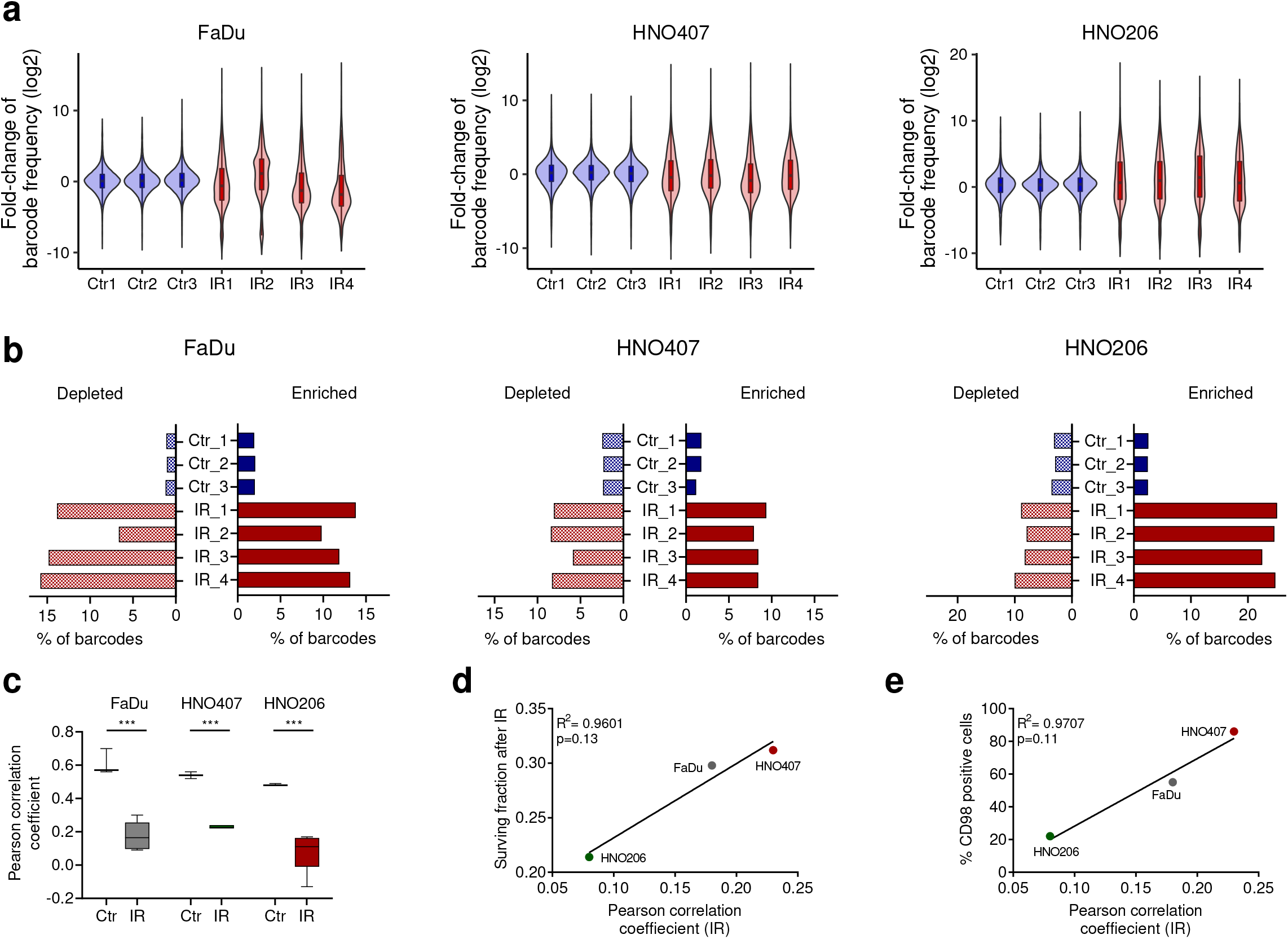
(**a**) Violin- and boxplots showing the fold-change of barcode frequency compared to initial starting population in each non-irradiated control (Ctr, blue) and irradiated (IR, red) replicate. The center line of the boxplots is the median, lower box bound is the 25% percentile, upper box bound is the 75% percentile, and whiskers are minimum and maximum values. (**b**) Bar plots show the proportion of enriched (≥4-fold (log_2_), red) and depleted (≤4-fold (log_2_), blue) barcodes in each non-irradiated (Ctr, blue) and irradiated (IR, red) replicate after normalization to the respective initial starting population. (**c**) Boxplots showing the Pearson correlation coefficients after pairwise comparison of clonal composition of non-irradiated (n=3) and irradiated (n=4) replicates in all HNSCC cell lines (unpaired t-test, ***, p<0.0001). The center line of the boxplots is the median, lower box bound is the 25% percentile, upper box bound is the 75% percentile, and whiskers are minimum and maximum values. (**d**+**e**) Linear correlation analysis of the average Pearson correlation coefficients of irradiated FaDu, HNO407 and HNO206 samples and (**d**) the surviving fraction at 4Gy (R^2^ = 0.9601, p=0.13) (**e**) the proportion of CD98 positive cells (R^2^ = 0.9707, p=0.11).

**Supplementary Figure 6.**
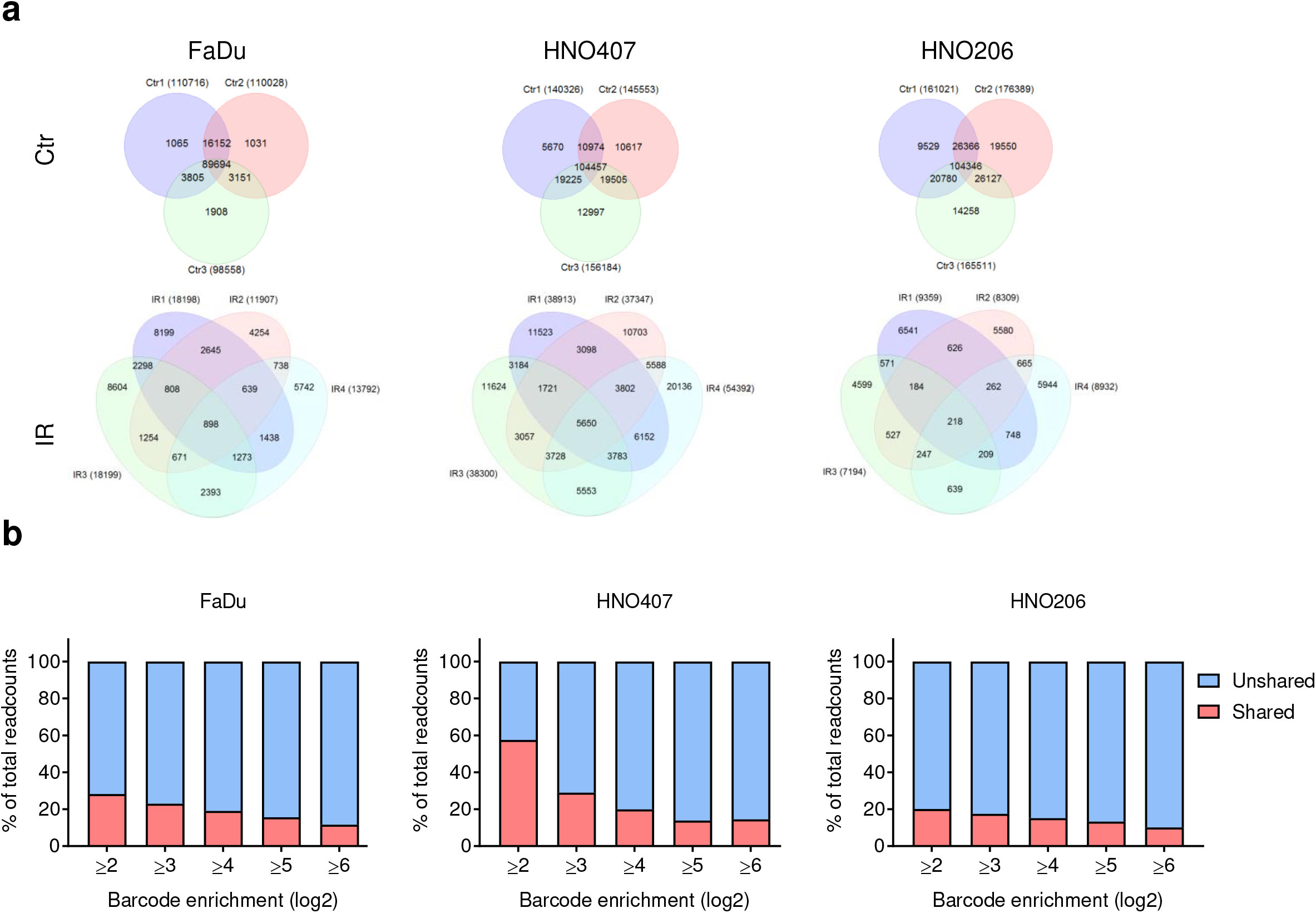
(**a**) Venn diagrams displaying the number of barcodes that are shared among the different non-irradiated control replicates (Ctr, n=3) and irradiated replicates (IR, n=4). The sum of barcodes that are detected in at least two replicates were calculated and defined as shared barcode population. In opposite, the sum of barcodes detected in only one replicate was defined as unshared barcodes. (**b**) Stacked bar plots showing the proportion of total sequencing readcounts derived from the deterministic cell population (shared barcodes, red) and stochastic cell population (unshared barcodes, blue) for different thresholds of barcode enrichment (log_2_).

**Supplementary Figure 7.**
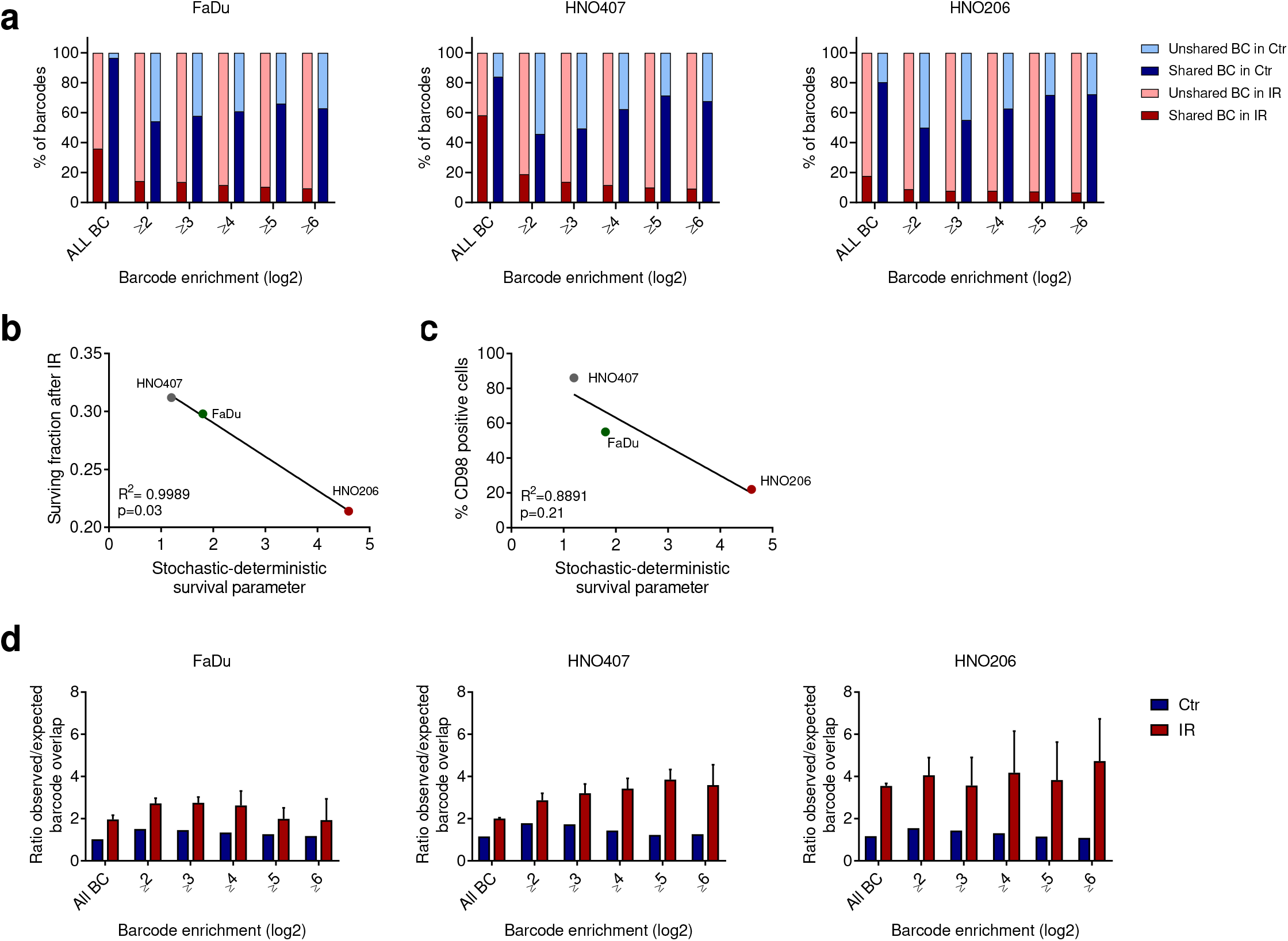
(**a**) Stacked bar plot showing the proportion of unshared barcodes and barcodes shared in at least two replicates in non-irradiated control (blue) and irradiated samples (red) in FaDu, HNO407 and HNO206 at different thresholds of barcode enrichment (log_2_). The proportions of shared and unshared barcodes were used to determine the stochastic-deterministic survival ratio. (**b+c**) Linear correlation analysis of the stochastic-deterministic survival ratio, which is calculated based on the proportion of non-shared clones divided by the proportion of shared clones, and (**b**) the surviving fraction of FaDu, HNO407 and HNO206 at 4Gy (R^2^=0.9989, p=0.03) and (**c**) the proportion of CD98 positive cells in FaDu, HNO407 and HNO206 (R^2^=0.8891, p=0.21). (**d**) Ratio of observed and expected barcode overlap based on a Fisher’s exact test in non-irradiated (Ctr, blue) and irradiated (IR, red) samples at different thresholds of barcode abundancy.

**Supplementary Table 1.**
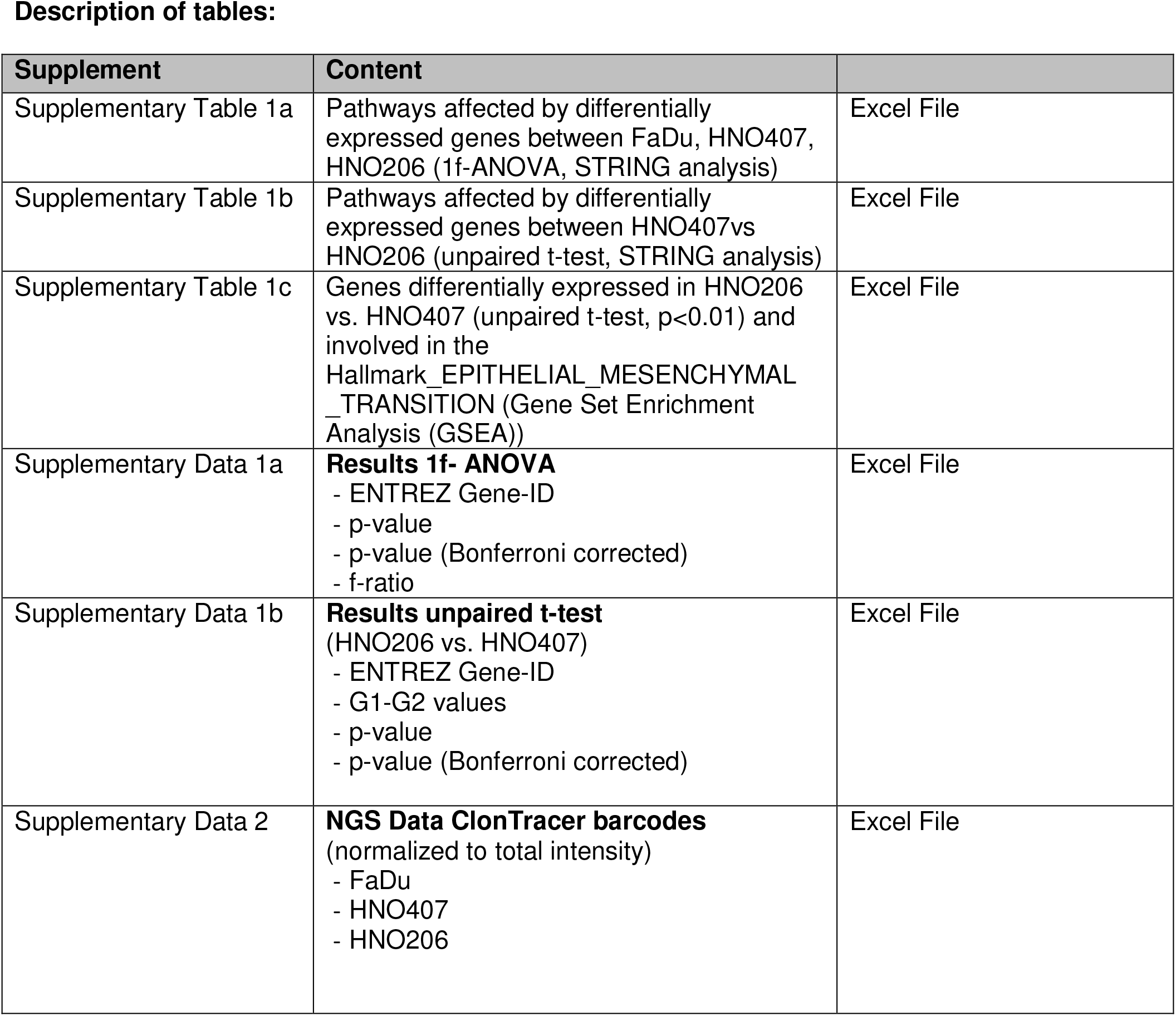
(**a**) Pathways and biological processes differentially regulated in FaDu, HNO407 and HNO206 (1-factor ANOVA, p<0.00005, FDR<0.05). Pathways affected by differentially expressed genes were identified using the Reactome pathway database. (**b**) Pathways affected by genes that are >4-fold differentially expressed in the patient-derived cell lines HNO206 (primary tumor) compared to HNO407 (lymph node metastasis), (unpaired t-test, p<0.001, log_2_, FDR<0.05). Pathways were identified using the Reactome database. (**c**) Genes that are differentially expressed in HNO206 compared to HNO407 (unpaired t-test, p<0.01) and involved in the Hallmarks gene set, which are differentially regulated in the HNO407 and HNO206 cell lines based on Gene Set Enrichment Analysis (GSEA). The Hallmark gene set collection from the Molecular Signature Database (MSigDB) was used for GSEA.

